# Cancer target discovery enabled by transcriptome-based virtual CRISPR screening

**DOI:** 10.1101/2024.10.24.620074

**Authors:** Ananthan Sadagopan, Bingchen Li, Jiao Li, Yantong Cui, Riva Deodhar, Di Yang, Yuqianxun Wu, Prathyusha Konda, Christy Biji, Dharma R. Thapa, Meha Thakur, Cary N. Weiss, Toni K. Choueiri, Jaimie H. Cheah, John G. Doench, Benjamin J. Drapkin, Srinivas R. Viswanathan

## Abstract

Functional genetic screens have uncovered dependencies in many cancers, but experimentally screened models for most cancers are far outnumbered by molecularly-profiled tumors, particularly for rare cancers. We used machine learning to infer gene dependencies from tumor transcriptional profiles, applying our model to the TCGA (11,373 tumors; 28 lineages), rare cancers (1,034 tumors, including 17 kidney cancer subtypes), and 509 previously unscreened cancer cell lines. Besides recovering dependencies previously identified in functional screens, we inferred drug response and synthetic essential relationships directly from tumors, including those associated with *RB1* inactivation, *KRAS* mutation, and microsatellite instability. Via dependency prediction, we discovered and validated a shared reliance on oxidative phosphorylation in two previously unscreened rare cancers both driven by *TFE3* gene fusions. We also nominate additional actionable vulnerabilities across various rare kidney cancers lacking experimental models. Our approach enables discovery of cancer vulnerabilities from transcriptomes, even in the absence of functional screening.

## Introduction

A cornerstone of precision oncology is matching therapies to specific cancers based on predictive molecular features. Recent advances in the molecular classification of cancer coupled with advances in genome-scale functional genetic screening have enabled the discovery of multiple biomarker-dependency pairs. Several of these dependencies have already been successfully translated into effective targeted therapies in specific cancers, with others in clinical trials (*1–9*).

However, two gaps continue to limit the broader application of this paradigm. First, many rare cancers have been poorly represented in large-scale functional genetics efforts due to a lack of robust cellular models, despite often having homogeneous genomic landscapes with singular driver alterations that are directly linked to robust vulnerabilities (*10–12*). Second, even common cancers may lack experimental models that reflect the heterogeneity observed in tumors. For example, prostate cancer, the most common cancer in men, has only ten cell lines included in the Cancer Dependency Map (DepMap), of which only three display dependence on the androgen receptor (*AR*) – the quintessential driver of nearly all prostate adenocarcinomas (*13*, *14*). Additionally, only one of these lines harbors the *TMPRSS2-ERG* gene fusion, which is found in nearly half of prostate tumors overall (*15*, *16*). As a result, dependencies inferred from functional screens on existing cell lines may not necessarily translate to an individual patient’s tumor.

Kidney cancer encapsulates both of the above challenges as it comprises dozens of distinct subtypes that vary in prevalence (*17*). Several rare subtypes of renal cell carcinoma (RCC) are not represented by experimental models while models of common subtypes only partially recapitulate the spectrum of genomic alterations found in corresponding tumors (*18*). These gaps have limited the ability to test the hypothesis that these distinct subtypes of kidney cancer vary in their dependency profiles. This is of clinical significance, since therapies designed for the most common kidney cancer (clear cell renal cell carcinoma, ccRCC) are frequently applied to other kidney cancers due to a lack of alternatives, yet they often yield lower response rates due to their distinct biology (*19–24*).

An exemplar rare kidney cancer is translocation renal cell carcinoma (tRCC) - driven by an activating gene fusion involving an MiT/TFE family transcription factor, most commonly *TFE3*. tRCC has a distinct genomic and transcriptional profile compared to other kidney cancers and lacks effective molecular targets (*25–30*). Furthermore, only a few cell line models of tRCC have been reported (*31–34*) and none were included in prior large scale screening efforts (*7, 35–37*), leading to some candidate-based vulnerabilities (*38–41*), but limiting unbiased functional screens that have been used for target discovery in more common cancers (*11*, *12*). In addition, *TFE3* fusions can drive a spectrum of other rare cancers apart from tRCC, including alveolar soft part sarcoma (ASPS) (*42*), perivascular epithelioid cell tumor (PEComa) (*43*), epithelioid haemangioendothelioma (EHE) (*44*), malignant chondroid syringoma (*45*), and ossifying fibromyxoid tumors (*46*). It remains unknown whether these tumor types, most of which do not have experimental models amenable to large-scale screening, have distinct dependency profiles from tRCC despite sharing the same driver fusion.

In this study, we applied machine learning to predict genetic dependencies based on tumor or cell line transcriptome profiles. We apply our predictive modeling to conduct “virtual” CRISPR screens across a broad range of cancer types – including those without experimental models previously subjected to functional CRISPR screening. Via this approach, we discover and validate several novel dependencies and synthetic essential relationships directly from tumor transcriptome profile.

## Results

### Establishing gene models for dependency prediction in tumor samples

We sought to establish machine-learning models that identify genetic dependencies from cell line or tumor genomic/transcriptomic data. A broader goal was to uncover vulnerabilities overlooked in prior functional screens and to nominate candidate dependencies in rare cancers without available cell line models.

We applied 5-fold cross-validation during a train-test cycle on the DepMap data set, in which >1100 cell lines experimentally subjected to CRISPR screening were also profiled by RNA-Seq (transcriptome data) or exome sequencing (mutation data). We first used both expression and mutation features from each cell line to calculate a predicted dependency score for each of 16,845 genes, selecting the best performing of 60 potential models for each gene (**Fig. 1A, Fig. S1A; Methods**). We observed that models trained on solely transcriptome profile versus those including both transcriptome and mutation data were highly correlated in performance (R=0.987); notable outliers for better prediction with mutation data included *NRAS*, *KRAS*, and *BRAF*, though these were all were still well-predicted using transcriptome data alone (**Fig.1B**). We therefore elected to proceed with transcriptome-only dependency prediction in order to establish a streamlined workflow (which we hereafter term **<u>Tr</u>**anscriptional **<u>P</u>r**ediction of **<u>Let</u>**hality, “TrPLet”) that could be clinically translated.

**Fig. 1.**
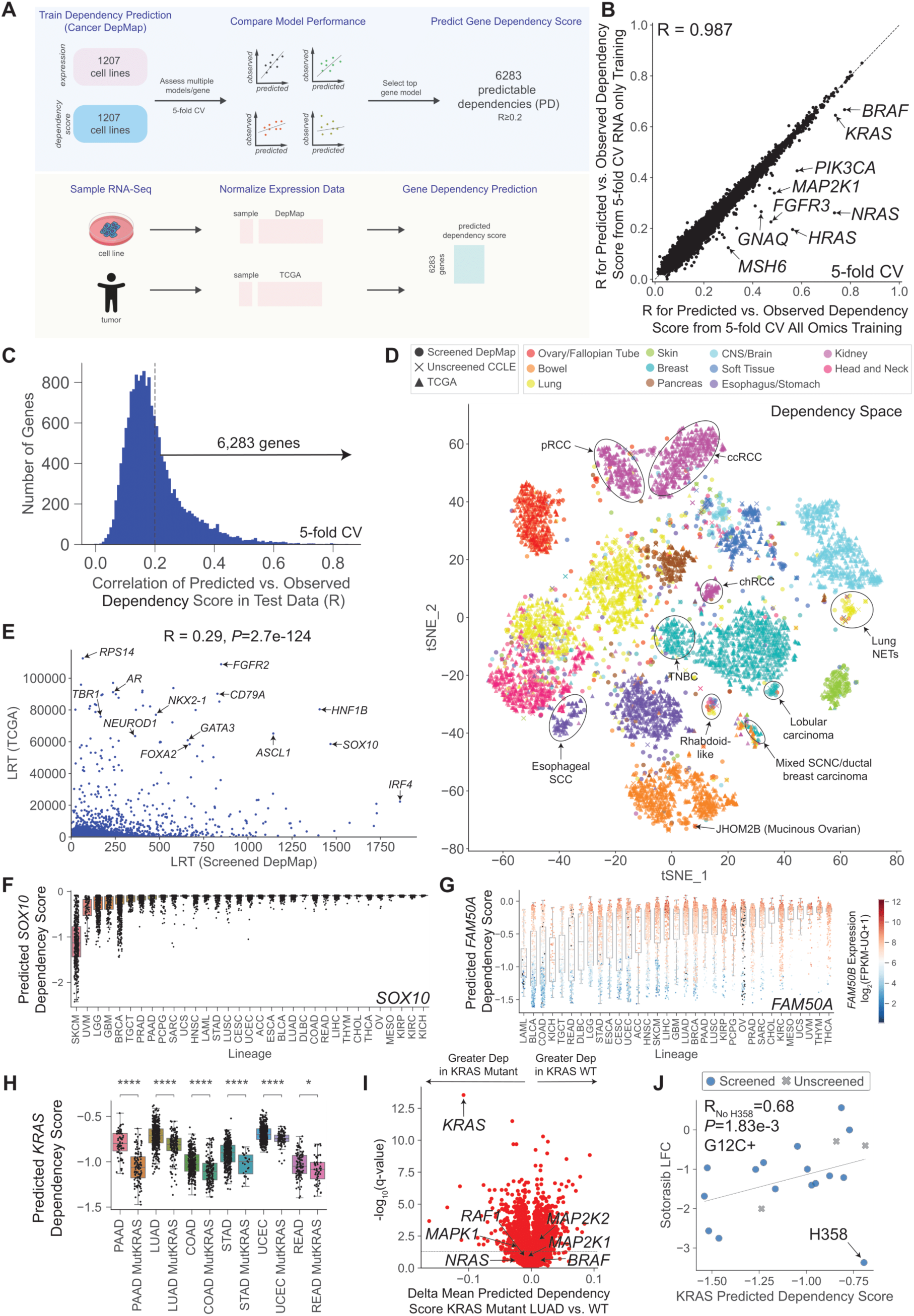
A machine-learning model predicts tumor and cell line dependencies from transcriptome profile. **(A)** Schematic of machine learning approach used to nominate candidate vulnerabilities in cell lines or tumors based on RNA-Seq profile. **(B)** Comparison of model performance for each PD gene using both RNA-seq and mutation features (x-axis) or RNA-seq only (y-axis). Performance for each model was assessed by computing the Pearson correlation coefficient between predicted dependency scores and screened Chronos scores for a given gene across DepMap (averaged from 5-fold cross-validation). Best model for each gene is shown. **(C)** Distribution of model performance for dependency prediction across all genes; best model for each gene is shown. 6,283 genes are highly predicable (“PD genes”) with an R ≥ 0.2 (threshold used for this study). **(D)** *t*-SNE projection based on dependency score for PD genes for tumors in TCGA predicted by our model (N=6,498, triangle), cell lines unscreened in CCLE predicted by our model (N=265, cross), and cell lines experimentally screened in the DepMap (N=1010, circle), restricted to common lineages. Tumors are colored based on annotated lineage and selected clusters are labeled. **(E)** Likelihood ratio test for skew-t vs. normal distribution for the distribution of each PD gene’s dependency score predicted in TCGA (y-axis) or screened in DepMap (x-axis). Selected outliers are labeled. **(F)** Predicted dependency scores for *SOX10* across TCGA tumor types. **(G)** Predicted dependency scores for *FAM50A* across tumor types profiled in TCGA. Samples are colored by *FAM50B* expression level. **(H)** Predicted *KRAS* dependency scores across the six lineages in TCGA with the greatest *KRAS* mutation frequency; tumors grouped by lineage and *KRAS* mutation status. *P*-values computed by Mann Whitney U-test; *P<0.05, ****P<0.0001. **(I)** Volcano plot of predicted dependencies across 6,283 PD genes in KRAS mutant vs. wild-type TCGA lung adenocarcinomas (LUAD). X-axis shows the difference in predicted dependency score between categories (stronger dependency in KRAS-mutant tumors on the left), y-axis shows the -log_10_(q-value) from BH-corrected heteroscedastic t-test. **(J)** Correlation between predicted *KRAS* dependency score and log_2_(fold-change) in cell viability upon single dose treatment (2.5 μM, 5 days) with the KRAS G12C inhibitor sotorasib in DepMap across KRAS^G12C^+ cancer cell lines. Pearson correlation coefficient and *P*-value computed from *t*-distribution are annotated; outlier cell line H358 excluded during calculations.

Transcriptome-only dependency prediction models were trained for 16,845 genes, using the best model for each gene (**Fig. S1A**). Of these, 6,283 genes were predictable at a performance threshold used in similar studies (*47*) (R ≥ 0.2 between experimentally derived Chronos dependency score and predicted dependency score). These 6,283 genes were hereafter termed <u>p</u>redictable <u>d</u>ependencies (“PD”) (**Fig. 1C, Fig. S1B, Table S1**). Notably, the number of PD genes significantly exceeds that in prior studies utilizing multiomic features to identify gene dependencies, potentially due to a larger training set and the selection of optimal prediction models on a per-gene basis (*47*, *48*) (**Fig. S1C-D**).

### Transcriptome-informed dependency prediction across cancer types

While prior efforts have sought to predict dependency profiles from multiomic data, these have typically predicted only a small number of genes with good accuracy owing to smaller training datasets, have had variable performance in tumor samples, or have not been applied to wholly new/unseen tumor types, such as previously uncharacterized rare cancers (*47–51*).

We first applied TrPLet – which reliably predicts dependency scores for ∼37% of widely expressed genes – to 11,373 tumors representing 33 lineages in The Cancer Genome Atlas (TCGA) (**Table S2**). When clustered on the basis of predicted dependency score (t-distributed Stochastic Neighbor Embedding, t-SNE), TCGA tumors clearly segregated by lineage with cell lines that were experimentally screened in the DepMap; by contrast, this was much less pronounced using RNA expression-based clustering (**Fig.1D**, circle vs. triangle**; Fig. S1E**).

We also predicted dependencies for 509 cancer cell lines with transcriptome data in the CCLE but not yet screened, revealing that predicted dependency profiles closely align with experimentally determined profiles within the same lineage (**Fig. 1D**, cross vs. circle; **Table S2**). Some informative exceptions included: (**1**) a mucinous-histology ovarian cell line (JHOM2B) clustering with bowel tumors/cell lines dependent on *KRAS, SOX9*, and Wnt signaling; treatment of mucinous ovarian cancer with gastrointestinal-type chemotherapy regimens is notably associated with better outcomes compared to gynecologic regimens (*52*). (**2**) lung neuroendocrine carcinomas clustering near CNS tumors rather than lung adenocarcinomas or squamous cell carcinomas, and (**3**) esophageal squamous carcinomas aligning with head and neck tumors instead of gastric adenocarcinomas. These findings highlight that a tumor’s dependency profile may be uncoupled from simply its organ of origin and suggest an approach to select cell line models suited to a clinical context of interest.

Moreover, experimentally determined selective dependencies in the DepMap correlated with predicted selective dependencies in TCGA (R=0.29, *P*=2.7e-124), with certain lineage dependencies such as *AR* (prostate), *TBR1* (brain), and *NKX2-1* (lung) being predicted with greater selectivity in TCGA than DepMap (**Fig. 1E**). This was unrelated to tumor purity (**Fig. S1F**) and may represent differences in cohort composition (e.g. percentage of prostate cell lines in DepMap vs. percentage of prostate tumors in TCGA) and/or differential prevalence of dependency-associated factors *in vivo* vs. *in vitro*.

We robustly recovered dependencies of various classes, including lineage-dependencies, paralog dependencies, and alteration-associated dependencies. For example, predicted dependency on *SOX10*, a melanocyte lineage gene (*53*), was the strongest in both cutaneous and uveal melanoma (*54*, *55*) while b-catenin (encoded by *CTNNB1*) was predicted to be the strongest dependency in colorectal cancers, consistent with prior studies (*56–58*) (**Fig. 1F**, **Fig. S1G**). Highly weighted expression features were biologically plausible in both cases, including *SOX10* and its target *TRIM51* (*59*) as predictors of *SOX10* dependency and *AXIN2*, *NKD1*, *ASCL2*, and *BMP4* as predictors of *CTNNB1* dependency (*60–62*). Predicted dependencies for *CDK4* also showed a lineage-skewing towards breast cancer, where CDK4/6 inhibitors are clinically approved (*63*) (**Fig. S1G-H**).

Paralog dependencies were also readily recovered: for instance, *FAM50A* dependency was predicted in tumors expressing low levels of *FAM50B* within and across lineages. These two paralogs of unknown function have been previously reported to be synthetically essential with each other via combinatorial CRISPR screening (*64*) (**Fig. 1G, Fig. S1H**). Additionally, dependencies on somatically altered driver genes were strongly predictable, even using only transcriptome data. For example, *ERG* dependency was predicted in prostate tumors harboring both *TMPRSS2-ERG* and *SLC45A3-ERG* fusions, despite only a single *ERG* fusion-positive prostate cell line in the DepMap training data (VCaP, **Fig. S1I**).

Similarly, *KRAS* dependency was the top predicted dependency in *KRAS*-mutant lung cancers, but other MAPK pathway members were not predicted to be selectively dependent in a *KRAS*-mutant context, despite having overlapping transcriptome signatures (*65*) (**Fig. 1H-I, Fig. S1H**). Moreover, predicted *KRAS* dependency score strongly correlated with the level of sotorasib sensitivity in cancer cell lines harboring a KRAS^G12C^ mutation (*66*) (R=0.68, *P*=1.83e-3, excluding outlier H358); there was no correlation in cell lines with other activating KRAS mutations (R=-0.101, *P*=0.234, **Fig. 1J, Fig. S1J**). The seven KRAS^G12C^ cell lines predicted to be most *KRAS* dependent (top 39%, analogous to the ∼40% clinical response rate to sotorasib (*67*, *68*)) displayed a significantly higher drug sensitivity to sotorasib than the remaining cell lines (*P*=9.5e-3). Although this bears further validation in clinically annotated patient datasets, it suggests that a transcriptome-informed, predicted *KRAS* dependency score can identify the subset of G12C mutant tumors that may respond best to sotorasib. We also observed that the correlation between sotorasib sensitivity in KRAS^G12C^+ cell lines and experimentally derived *KRAS* dependency score (Chronos score, R=0.216, *P*=0.423) was substantially lower than when using the predicted *KRAS* dependency score; this perhaps reflects the heterogeneous experimental factors influencing CRISPR-mediated knockout phenotypes (**Fig. S1J**).

Beyond sotorasib, sensitivity to several other compounds correlated strongly with predicted dependency scores on their known targets, including MDM2 inhibitors (idasanutlin, AMG-232), BCL2 inhibitors (venetoclax, navitoclax), BRAF inhibitors (dabrafenib, PLX-4720, SB-590885), an EGFR inhibitor (afatinib), a HER2 inhibitor (CP-724714), a FGFR2 inhibitor (AZD4547), an IGF1R inhibitor (BMS-754807), a PIK3CD inhibitor (idelalisib), and a GPX4 inhibitor (ML210) (**Fig. S1K**). Notably, several genes displayed greater variance in predicted dependency score across TCGA tumors versus normal tissues (GTEx), suggestive of a favorable cancer-specific therapeutic window to their targeting (**Fig. S1L**).

Therefore, our TrPLet pipeline can identify various classes of biologically meaningful genetic dependencies (many of which are linked to the sensitivity of specific compounds) and may also be able to nominate transcriptomic biomarkers of response to these targeted therapies.

### Discovery of Synthetic Lethal Relationships in the TCGA

As the TCGA cohort is nearly 10-fold larger than DepMap (n=11,373 vs. n=1,207) and carries a different spectrum of somatic alterations, we reasoned that our dependency prediction approach could recover synthetic lethal relationships for which DepMap may be underpowered. We therefore used our approach to conduct “*virtual*” synthetic essential CRISPR screening across the TCGA, using dependency scores for PD genes to identify candidate synthetic essential relationships with altered cancer genome census (CGC) genes. For each CGC gene (n=753), we compared dependency scores for every PD gene in CGC gene-altered vs. unaltered samples (**Fig. 2A**). Alterations by mutation and deletion were considered separately, and the same analysis was also performed on the DepMap using experimentally-derived Chronos dependency scores.

**Fig. 2.**
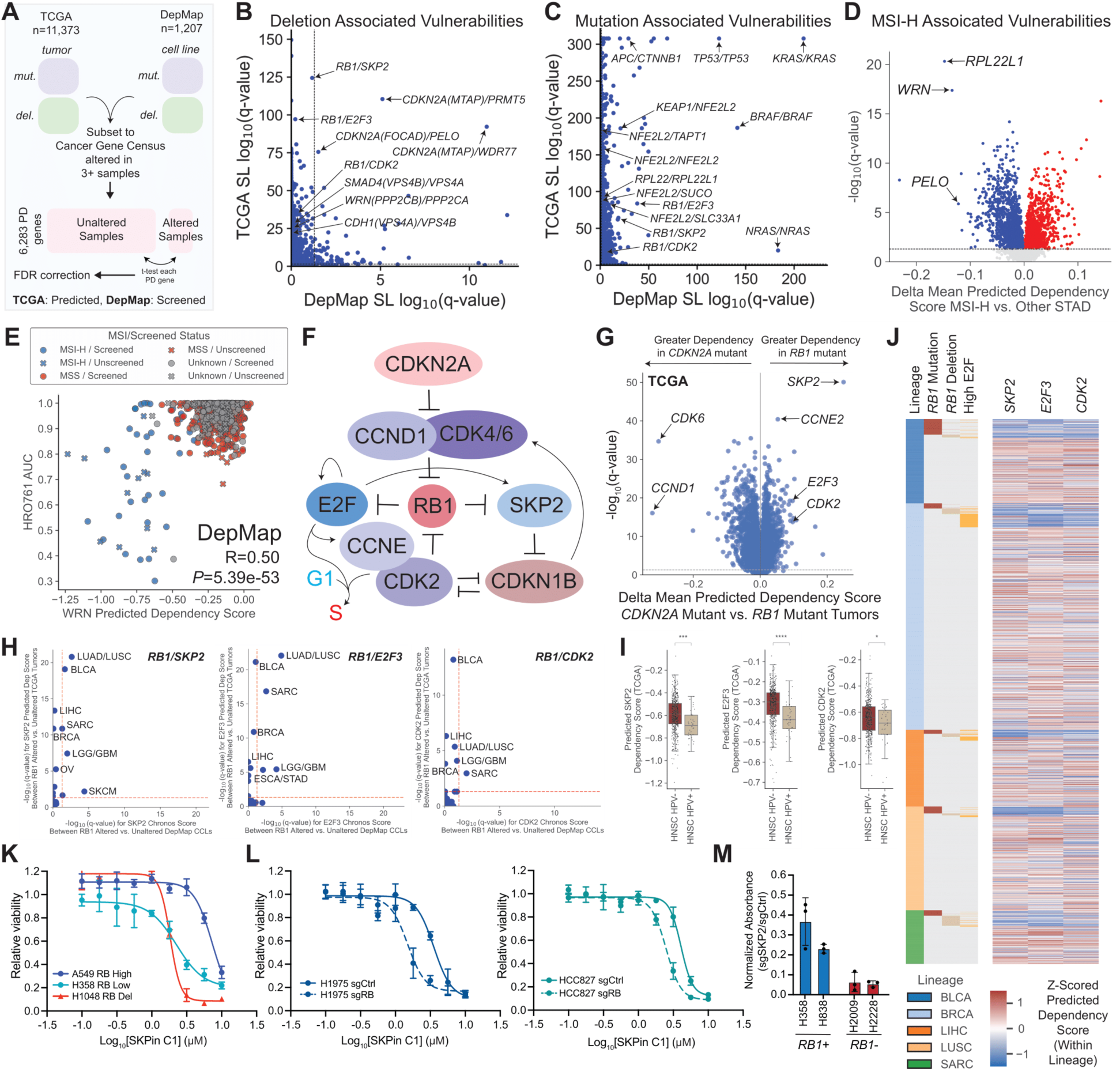
Synthetic lethal inference across TCGA via virtual CRISPR screening identifies SKP2 dependency in Rb1-altered cancer cells. **(A)** Strategy for synthetic lethal discovery across the DepMap and TCGA, for all Cancer Genome Census (CGC) genes altered in 3 or more samples within each cohort (N=753). Deletions and mutations were analyzed separately. **(B-C)** Comparison of -log_10_(q-values) for synthetic lethal pairs with CGC genes predicted in TCGA (y-axis) or screened in DepMap (x-axis). Lines at q=0.05 are shown on the plot. If an alteration was represented in TCGA but not DepMap, the q-value of the synthetic lethal pair was set to 1 in DepMap. Separate plots are shown for CGC genes altered by: **(B)** deletion, with relevant collaterally deleted genes shown in parentheses and **(C)** mutation. **(D)** Comparison of predicted dependency scores in PD genes between MSI-H and MSS TCGA stomach adenocarcinomas (STAD). X-axis shows the difference in predicted dependency score, y-axis shows the -log_10_(q-value) from BH-corrected heteroscedastic t-test. **(E)** Correlation between predicted *WRN* dependency score and sensitivity to the WRN inhibitor HRO761 profiled in the DepMap. Pearson correlation coefficient and *P*-value computed from t-distribution are annotated. **(F)** Schematic of Rb pathway, including known genetic relationships of *SKP2*, *CDK2*, and *E2F3*. **(G)** Volcano plot of predicted dependencies across 6,283 PD genes in *RB1* mutant vs. *CDKN2A* mutant TCGA samples. X-axis shows the difference in predicted dependency score between groups, y-axis shows the -log_10_(q-value) from BH-corrected heteroscedastic t-test. **(H)** -log_10_(q-values) from pairwise comparisons of *SKP2*, *E2F3*, or *CDK2* dependency score (predicted for TCGA, screened Chronos scores for DepMap) between *RB1* altered vs. unaltered samples within a lineage with Mann Whitney U-test. TCGA is shown on the y-axis, DepMap is shown on the x-axis; each point represents a lineage shared between TCGA and DepMap. **(I)** Predicted dependency scores for *SKP2, E2F3*, and *CDK2* across TCGA HNSC tumors by HPV infection status. *P*-values computed by Mann Whitney U-test; *P<0.05, ***P<0.001, ****P<0.0001. **(J)** Heatmap of predicted dependency scores for *SKP2, E2F3*, and *CDK2* (Z-scored within lineage) across lineages with frequent Rb alterations, grouped by RB1 mutation, RB1 deletion, or high E2F ssGSEA score (Z ≥1.5 within lineage). **(K)** Lung cancer cell lines (RB1 high: A549, RB1 low: H358, RB1 deleted: H1048; as assessed by Western blot in **Fig. S3D**) treated with indicated concentrations of SKPin C1 (SKP2 inhibitor) and assayed for cell viability after 3 days with CellTiter-Glo. Viability at each concentration is relative to vehicle-treated cells, shown as mean +/- s.d., n=3/4 biological replicates. **(L)** Rb-proficient NSCLC cell lines (H1975, HCC827) after CRISPR/Cas9 knockout of *RB1* or non-targeting sgRNA treated with indicated concentrations of SKPin C1 (SKP2 inhibitor) and assayed for cell viability after 3 days with CellTiter-Glo. Viability at each concentration is relative to vehicle-treated cells, shown as mean +/- s.d., n=3/4 biological replicates. **(M)** Quantification of colony formation assay for RB1+ (H358, H858) and RB1-(H2009, H2228) NSCLC cell lines after CRISPR/Cas9 knockout of *SKP2* (normalized to sgCtrl). Shown as mean +/- s.d., n=3 biological replicates per condition.

The strongest deletion-associated synthetic lethal relationships were *CDKN2A^Δ^*/*PRMT5* and *CDKN2A^Δ^*/*WDR77*, identified in both our DepMap and TCGA analyses (**Fig. 2B**). These represent collateral dependencies (*69*) to the queried CGC gene (*CDKN2A*) and are due to co-deletion of an adjacent gene (*MTAP*), leading to dependency on the WDR77-dependent methylosome; this relationship was originally elucidated via the DepMap (*70*, *71*). Similarly, our approach also identified *PELO* as a synthetic lethality associated with *CDKN2A* deletion, another collateral relationship recently shown to be due to biallelic co-deletion of the 9p21.3 gene *FOCAD*, a stabilizer of the superkiller complex (SKIc) (*72*).

Numerous additional dependencies validated in recent studies were readily identified via our TCGA dependency prediction, including paralog dependency pairs (e.g. *VPS4A/4B* (*73*), *RPL22*/*RPL22L1* (*74*, *75*)) and synthetic lethal interactions with the NRF2 pathway (e.g. *TAPT1*, *SUCO*, *SLC33A1* (*76*) and *KEAP1^mut^/NFE2L2* (*77*)) and *APC* (*CTNNB1* (*78*)) (**Fig. 2B-C**). Beyond dependencies associated with single gene alterations, we also identified vulnerabilities associated with microsatellite instability-high tumors (MSI-H): our *in silico* approach recovered all recently reported MSI-H-associated dependencies including *WRN* (*79*, *80*), *PELO* (*72*), and *RPL22L1* (*75*) (**Fig. 2D, Fig. S2A-C**), with biologically relevant predictive features in each model (**Fig. S2D**). Moreover, predicted *WRN* dependency score correlated with sensitivity to the small molecule WRN inhibitor: HRO761 across DepMap (R=0.50, *P*=5.39e-53, **Fig. 2E**).

In addition, our TCGA analysis revealed a previously uncharacterized collateral dependency associated with *WRN* deletion, and attributable to co-deletion of the nearby gene *PPP2CB* (deep deletion seen in ∼1.5% of tumors across the TCGA but nearly absent in the DepMap, **Fig. 2B, Fig S2E**). Heterozygous deletion of *PPP2CB*, which most often occurs in the setting of chr8p loss (*81*), was predicted to unveil dependency on the paralog gene, *PPP2CA*, a relationship suggested in two recent studies of paralog lethality (*82*, *83*). These paralogs share ∼97% protein homology (*84*) and encode the catalytic subunit of the PP2A phosphatase (an essential serine threonine phosphatase with diverse roles in regulating cell signaling (*85*)). Via comparison of single or double knockout of *PPP2CA/PPP2CB* in wild type and *PPP2CB*-deleted cell lines, we observed that the latter were more sensitive to double knockout; differential sensitivity to double knockout rather than single knockout of *PPP2CA* likely represents a dosage-dependent phenotype in the setting of incomplete CRISPR knockout and potential compensation of *PPP2CB* expression in the setting of heterozygous deletions (**Fig. S2F-N**).

### Virtual CRISPR screening pinpoints multiple synthetic essential relationships with Rb inactivation

Given our facile recovery of a wide array of gene dependencies previously validated via functional approaches, we next sought to determine whether applying TrPLet to the large TCGA dataset could nominate additional dependencies not rising to significance in the DepMap. This approach prominently highlighted three potential dependencies associated with *RB1* alteration: *SKP2*, *E2F3*, and *CDK2*. While these genes are linked to the Rb pathway and have been implicated as candidate vulnerabilities in individual contexts (*86–92*), they do not reach significance in a pan-DepMap synthetic lethal analysis, possibly due to underrepresentation of *RB1* deleted cell line models in the DepMap compared to the TCGA and the diverse mechanisms that converge on *RB1* inactivation (**Fig. 2B-C**, **Fig. S2E**).

To further explore this finding, we first compared predicted dependency scores between *RB1* mutant cancers (which lack expression of functional retinoblastoma protein, pRB) and *CDKN2A* mutant cancers (in which loss of p16 expression post-translationally inactivates pRB through de-repression of Cyclin D-CDK4/6 complexes) (*93*). As expected, the Cyclin D*-*CDK4/6 complex was identified as a selective vulnerability in *CDKN2A* mutant cancers, and consistent with *RB1* alterations as predictors of resistance to CDK4/6 inhibitors (*94*). Strikingly, however, *SKP2, E2F3,* and *CDK2* scored comparably strongly as selective vulnerabilities of *RB1*-mutant cancers (**Fig. 2F-G**). Importantly, while these Rb1-associated synthetic lethals were recovered in both TCGA and the DepMap in individual lineages, *SKP2*, *E2F3*, and *CDK2* scored in many more lineages in the TCGA (predicted dependencies, 9-11 lineages) than in the DepMap (experimentally-derived, 1-5 lineages), illustrating the increased power in performing this type of analysis across the TCGA (**Fig. 2H, Fig. S3A-B**). We also observed that synthetic lethal associations with *SKP2, E2F3,* and *CDK2* extended beyond somatic alterations of *RB1* to include oncoprotein-driven inactivation of pRB, such as in human papilloma virus (HPV)+ head and neck cancers in which pRB is bound by the HPV E7 oncoprotein (*95*) (**Fig. 2I**).

Inactivation of pRB promotes the G1-S transition via multiple effector pathways that involve *SKP2*, *E2F3*, and *CDK2* (**Fig. 2F**). Activation of E2F proteins leads to Cyclin E transactivation and activation of the Cyclin E-CDK2 complex (*96*, *97*). E2Fs also activate SKP2 transcription, leading to degradation of CDKN1B (p27), which relieves inhibition on Cyclin E-CDK2 (*98–103*). These relationships are reinforced by many additional feedback circuits, including mutual inhibition between Cyclin E-CDK2 and p27, positive feedback of E2Fs on their own transcription, inhibition of SKP2 by the C-terminal domain of pRB, and transactivation of DNA synthesis genes by E2Fs to promote the G1-S transition directly (*99*, *104–111*). Consistent with these complex, interconnected relationships, the predictive features of dependencies on *SKP2, E2F3*, and *CDK2* were overlapping and all enriched for *RB1* and pRB pathway genes (**Fig. S3C**). Moreover, there was strong predicted co-dependency between all three genes, which were enriched both in samples containing genetic *RB1* inactivation or in those with an elevated E2F transcriptional signature (**Fig. 2J**).

While *CDK2*, *SKP2* and *E2F3* vulnerabilities have been linked to Rb inactivation in prior studies, most have been limited to one or a few models (*89–91*, *112–115*), with a heavy emphasis on retinoblastoma and small cell lung cancer (SCLC), where biallelic genomic loss of *RB1* is nearly universal (*116–120*). Notably, however, SCLC is not represented in the TCGA dataset, from which these dependencies were originally nominated in our study. Given our observation that these genes may represent synthetic essentialities with *RB1* inactivation across diverse tumor types (**Fig. 2H**), we next sought to experimentally validate the window for SKP2 inhibition (the strongest scoring amongst the candidate Rb synthetic lethals). Using either CRISPR-mediated knockout or a recently developed small molecule inhibitor (SKPin C1 (*87*, *121*)) across a panel of Rb-deficient and Rb-proficient lung cancer cell lines (both SCLC and non-small cell lung cancer, NSCLC), we observed substantially increased sensitivity to SKP2 inhibition/genetic knockout in Rb-deficient versus Rb-proficient lines (**Fig. 2K-M, Fig. S3D-F**). Moreover, increased sensitivity to SKPin C1 was observed upon *RB1* deletion in two pairs of isogenic NSCLC lines in which *RB1* was deleted via CRISPR (**Fig. 2L**).

Overall, we conclude that our approach of dependency prediction – applied to a large cohort representative of the spectrum of tumors encountered in real world practice – can reliably nominate vulnerabilities and recover biologically meaningful synthetic essential relationships, even in contexts where functional screens may be underpowered for discovery.

### Nominating vulnerabilities in rare kidney cancers

Having established the tractability of our approach for nominating dependencies from tumor transcriptomes across the TCGA, we next turned our attention to kidney cancers, which comprise a notoriously heterogeneous group of > 40 molecularly distinct subtypes in both adults and children (*17*). Many of these cancers are rare, lacking in cell line models, and poorly represented or unrepresented in the TCGA – making them a prime application for dependency prediction.

We collected RNA-Seq data from 938 kidney tumors representing 17 subtypes across seven published studies (**Fig. 3A, Table S2**) (*122–130*). Using TrPLet, we then calculated predicted dependency scores for 6,283 PD genes and clustered tumors based on dependency profile, together with 22 renal cancer cell lines from the DepMap (on which dependency scores were experimentally determined) (**Fig. 3B**).

**Fig. 3.**
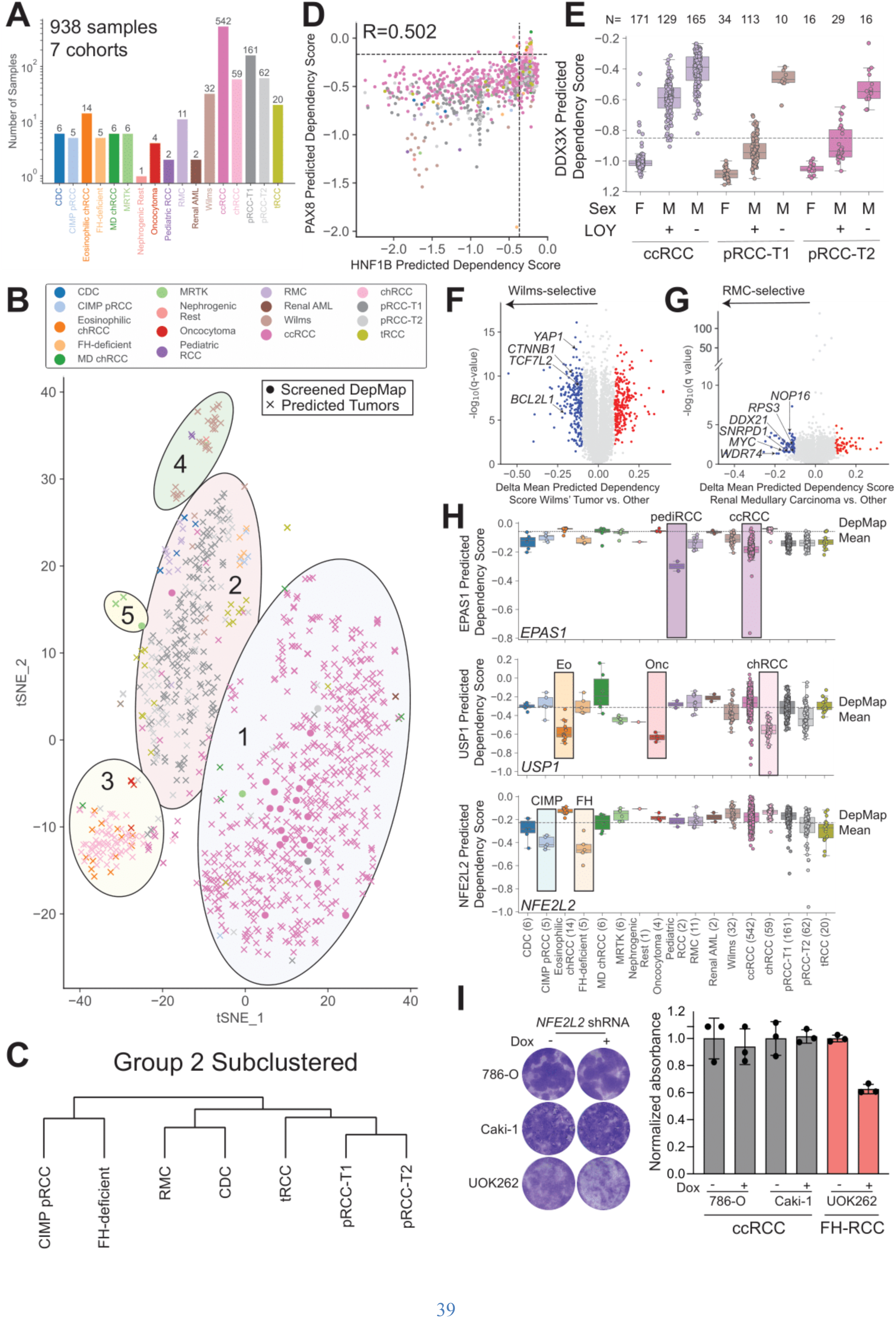
A landscape of candidate dependencies across rare kidney cancers. **(A)** RNA-Seq data from 938 renal tumor samples were curated from 7 published datasets, representing 17 distinct types of kidney cancer. Number of samples in each kidney cancer subtype is shown. **(B)** *t*-SNE projection based on dependency score for kidney tumors with dependencies predicted by our model (N=938, cross) and kidney cancer cell lines experimentally screened in the DepMap (N=22), across 6,283 PD genes (Z-scored to DepMap pan-cancer value for screened cell lines). Tumors are colored based on annotated subtype and five groups are outlined. **(C)** Dendrogram subclustering kidney cancer subtypes in group 2 based on dependency scores across 6,283 PD genes (hierarchical clustering, Z-scored to DepMap pan-cancer value for screened cell lines). **(D)** Scatter plot of *HNF1B* predicted dependency score versus *PAX8* predicted dependency score for individual kidney tumors (N=938), colored by subtype as indicated in Fig. 3B. Mean Chronos scores for *HNF1B* and *PAX8* in DepMap are indicated as dotted lines. Pearson correlation coefficient is shown. **(E)** Box plot of predicted *DDX3X* dependency score across ccRCC, pRCC-T1, and pRCC-T2 tumors in TCGA stratified by annotated sex and loss of chrY (LOY) status (*140*). Mean Chronos score for *DDX3X* in DepMap is indicated as a dotted line. **(F)** Volcano plot of differential dependencies between Wilms’ tumors and non-Wilms’ kidney tumors in the cohort. -log_10_(q-values) calculated from BH-corrected heteroscedastic t-test (y-axis) and difference in dependency scores between groups are shown (x-axis, blue: Δ<-0.1, q<0.05; red: Δ>0.1, q<0.05). **(G)** Volcano plot of differential dependencies between renal medullary carcinoma (RMC) tumors and non-RMC kidney tumors in the cohort. -log_10_(q-values) calculated from BH-corrected heteroscedastic t-test (y-axis) and difference in dependency scores between groups are shown (x-axis, blue: Δ<-0.1, q<0.05; red: Δ>0.1, q<0.05). **(H)** Box plot of predicted *NFE2L2* (top), *EPAS1* (middle), and *USP1* (bottom) dependency scores for individual kidney tumors (N=938), grouped by subtype indicated in Fig. 3A. Mean Chronos score in DepMap (experimentally derived) for these genes are indicated by dotted lines. **(I)** Representative image (left) and quantification of colony formation assay (right) for ccRCC (786-O, Caki-1) or FH-RCC cells (UOK262) after dox-inducible shRNA-mediated knockout of *NFE2L2*. Shown as mean +/- s.d., n=3 biological replicates per condition. *P*-values were calculated by heteroscedastic t-test compared to the no dox condition.

Kidney tumors and cancer cell lines collapsed into five broad dependency groups: (**1**) Group 1 was comprised primarily of ccRCC tumors and most RCC cell lines (which are enriched for clear-cell type (*131*)), as well as metabolically divergent chRCC (MD-chRCC) and renal angiomyolipoma (AML) tumors. MD-chRCC have been previously described as a distinct, clinically aggressive subset of chRCC with a distinct methylation and copy number alteration profile relative to classical chRCC (*124*); most MD-chRCCs also demonstrate sarcomatoid differentiation, which is observed in a subset of ccRCC tumors (*124*, *132*). (**2**) Group 2 comprised papillary RCC (both pRCC type 1 and type 2 as annotated in TCGA) as well as a number of diverse entities that were historically classified as pRCC type 2 (*133*), including CpG island methylator phenotype RCC (CIMP-RCC) and fumarate hydratase (FH)-deficient RCC. (**3**) Group 3 consisted of oncocytic tumors (oncoytoma, chRCC, eosinophilic chRCC). (**4**) Group 4 was comprised of Wilms’ tumors, nephrogenic rest (precursor to Wilms’ tumors), and pediatric RCCs which are rare pediatric tumors with distinct clinical characteristics and histopathology compared to adult RCCs (*134*). (**5**) Group 5 consisted of malignant rhabdoid tumors of the kidney (MRTK) and an MRTK cell line (**Fig. 3B**). Hierarchical clustering of dependency profiles of samples within group 2 elicited further sub-stratification of these diverse tumor types and separations based on RNA-based clustering alone were far less granular (**Fig. 3C, Fig. S4A**).

We then interrogated these data to identify differential dependencies across kidney cancer subtypes. The lineage transcription factors *HNF1B* and *PAX8* were predicted to be very strong dependencies in most RCC subtypes with the notable exception of the oncocytic tumors (chRCC, eosinophilic chRCC, oncoytoma, and MD chRCC) (**Fig. 3D**); this is consistent with these tumors arising from mitochondria-rich cells of the distal nephron (*135*) while other kidney cancers arise from proximal tubule kidney epithelial cells. Notably, *PAX8* and *HNF1B* vary in expression level throughout the developing kidney, with higher expression in the proximal pronephros (*136*), and these lineage dependencies are highly correlated to expression level (*137*, *138*).

*DDX3X*, a paralog dependency known to be unmasked by loss of the Y-chromosome encoded paralog *DDX3Y* (*139*), also varied across subtypes. *DDX3X* dependency was predicted to be pervasive in pRCC-T1, a tumor type with almost universal somatic loss of the Y chromosome (LOY) (*140*) and associated with LOY in other RCC subtypes as well (**Fig. 3E**). Similarly, dependency on *MDM2* was associated with *TP53* mutation status across RCC subtypes, with most kidney tumors expected to be dependent on *MDM2* in accordance with the low frequency of *TP53* mutations in kidney cancer (*141*) (**Fig. S4B**).

Predicted dependency on several selenium metabolism-related genes (*SEPHS2*, *SEPSECS*, *GPX4*, *PSTK*) suggested strong ferroptosis-associated cell death dependencies in ccRCC and renal AML, consistent with earlier studies in clear-cell tumors and a mesenchymal origin to renal AML (*142*, *143*). Notably, renal tumors with sarcomatoid and/or rhabdoid differentiation (S/R RCCs) were also predicted to share dependencies on selenium metabolism and antioxidant defense mechanism genes, while showing reduced reliance on lineage factors, supporting the model that S/R RCCs reflect a de-differentiation event that can occur in tumors of diverse parental histologies (*132*) (**Fig. S4C; Table S2**). Kidney cancer subtypes also exhibited varying predicted dependency on apoptosis-related genes, with pRCCs, Wilms’ tumors, nephrogenic rests, and pediatric RCCs showing particular reliance on *BCL2L1*, and most kidney cancers (except for renal AML and MD-chRCC) being relatively resistant to *MCL1* knockout **(Fig. S4D**). These findings suggest distinct subtype-specific vulnerabilities to cell death mechanisms across kidney cancers.

Our analysis also revealed dependencies highly enriched in one or two subtypes of kidney cancer. For example, Wilms’ tumors had predicted dependency on Wnt/β-catenin signaling (*CTNNB1*, *TCF7L2*), as well as on *YAP1* and *BCL2L1*, which have been shown to be essential for the survival of β-catenin-driven cancers (*144*); these dependencies are consistent with frequent Wnt/β-catenin alterations in Wilms’ tumor (*145–147*) (**Fig. 3F**). Additionally, renal medullary carcinoma (RMC) displayed many predicted dependencies in the Myc pathway, consistent with a recent report describing strong upregulation of the Myc transcriptional program in this lineage (*148*) (**Fig. 3G**).

We also recovered *EPAS1* (HIF-2α) dependency in ccRCC, consistent with activated hypoxia signaling secondary to near universal *VHL* deletion and the demonstrated clinical effectiveness of HIF-2α inhibitors in this subtype (*149*); notably, pediatric RCCs also displayed predicted *EPAS1* dependency despite clustering apart from adult ccRCC in dependency space, and despite the fact that *VHL* loss is uncommon in pediatric RCC (*150*), while tRCCs (which do not harbor *VHL* alterations (*24*)) were not predicted to be *EPAS1*-dependent (**Fig. 3H**). We identified ubiquitin-specific protease 1 (USP1) as a strong, selective, and druggable candidate dependency in chromophobe/oncocytic tumors, which lack both cell line and PDX experimental models for rigorous preclinical validation (**Fig. 3H**). Finally, we observed stark differences in predicted *NFE2L2* dependency across RCC subtypes; FH-deficient RCC and CIMP-RCC were predicted to be strongly dependent on *NFE2L2*, in accordance with prior reports of NRF2 pathway activation in these subtypes via methylation of *KEAP1* (CIMP-RCC) or succinylation of *KEAP1* (FH-RCC) (*151–153*). Consistent with our predictions, we validated that shRNA-mediated knockdown of *NFE2L2* was selectively deleterious in the FH-RCC cell line, UOK262 (*154*), which was not included in the DepMap (**Fig. 3I, Fig. S4E**).

Overall, we provide a landscape of candidate dependencies in rare kidney cancers that could be used as a starting point to develop mechanism-inspired therapeutic strategies in these diseases.

### Integrating predicted dependencies and functional screens to map vulnerabilities in tRCC

Translocation renal cell carcinoma is a prime example for this study’s overall rationale: (**1**) it is a rare subtype of kidney cancer with no effective treatments, and for which treatments used for other subtypes of RCC are less effective (*19–22*). (**2**) while a few tRCC cell line models exist, these are outnumbered by transcriptionally profiled tumors by approximately 10-fold. (**3**) no tRCC cell lines have yet been included in the DepMap, so the genome-wide dependency landscape in this cancer remains unknown.

We sought to identify potential dependencies in tRCC by applying our predictive model on tRCC tumor samples. We then integrated these findings with functional screens and experimental validation in currently available tRCC cell line models.

We first applied our model to calculate predicted dependency scores for 6,283 PD genes in 88 tRCC tumors and seven tRCC cell lines profiled by RNA-seq in either this or prior studies (**Table S2**). Using ccRCC as a lineage-matched control cancer, we nominated tRCC-selective dependencies by comparing predicted dependency scores for PD genes in tRCC cell lines/tumors to experimentally-derived dependency scores for the same genes amongst ccRCC cell lines in the DepMap.

This approach enabled us to nominate dependencies that vary both within and between lineages, which we validated genetically and/or chemically. For example, *MDM2*, which we nominated above as a dependency across several kidney cancer subtypes, was predicted to be a strong dependency in most tRCCs but displayed substantial variation within and across subtypes that tracked with *TP53* mutation status (**Fig. S5A-F**). Other genes were predicted to be strong dependencies in a subset of tRCCs but without clear biomarkers, such as *KMT2D* dependency in the s-TFE tRCC cell line (**Fig. S5G-I**).

By contrast, when comparing between subtypes (tRCC vs. ccRCC), we predicted uniform tRCC-selective dependency on several mitochondrial proteins (*ATP5F1D, GRPEL1, IARS2, LETM1, PTCD3, SOD2, YME1L1, UQCRFS1*) and hypoxia-inducible factor signaling (*VHL, EGLN1*) from tumors and cell lines (**Fig. 4A-B, Fig. S6A-C**). Ontology analysis of top tRCC-selective dependencies confirmed enrichment for genes involved in oxidative phosphorylation, mitochondrial translation, and electron transport (**Fig. 5C**), consistent with our recent metabolic study reporting heightened aerobic respiration in tRCC (*155*).

**Fig. 4.**
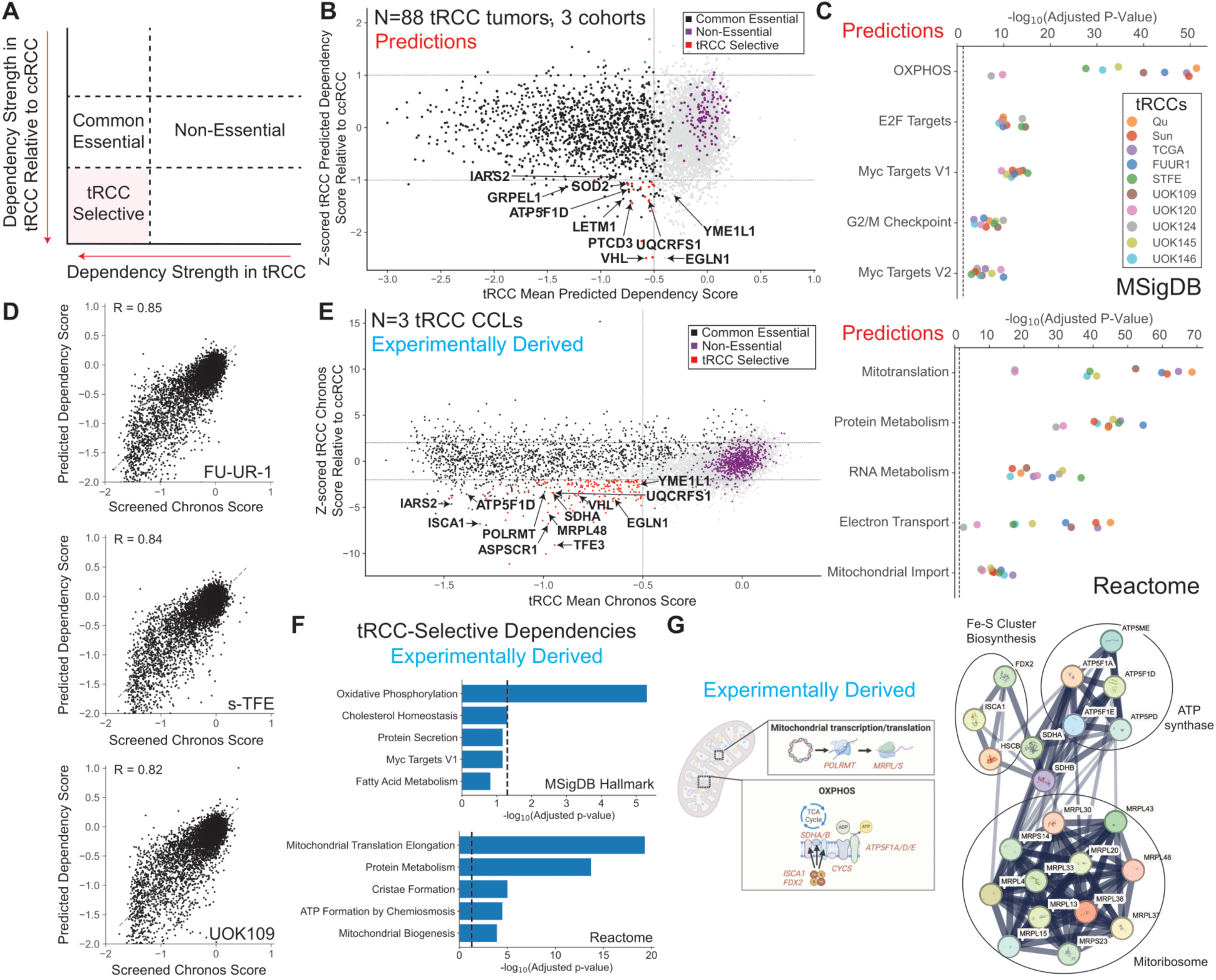
Virtual and experimental CRISPR screens converge on oxidative phosphorylation dependency in translocation renal cell carcinoma. **(A)** Schematic for identification of tRCC-selective genetic dependencies. **(B)** Mean predicted dependency score for each PD gene across 88 tRCC tumors from 3 tRCC cohorts plotted against the Z-scored predicted dependency score for that gene (relative to screened DepMap ccRCC cell lines). Non-essential genes are colored purple while common essential genes are colored black (*184*). tRCC-selective dependencies are colored in red. **(C)** Pathway enrichment (Enrichr) on predicted tRCC-selective dependencies (relative to ccRCC) in tRCC tumor cohorts (Qu, Sun, TCGA) and tRCC cell lines (FU-UR-1, S-TFE, UOK109, UOK120, UOK124, UOK145, UOK146). **(D)** Correlation between Chronos score from CRISPR screens performed in this study and dependency score predicted by our model for 6,283 PD genes across three tRCC cell lines (FU-UR-1, S-TFE, UOK109). Pearson correlation coefficient is shown for each cell line. **(E)** Mean Chronos score for each gene across three screened tRCC cell lines plotted against the Z-scored Chronos score for that gene (relative to screened DepMap ccRCC cell lines). Non-essential genes are colored purple while common essential genes are colored black (*184*). tRCC-selective dependencies are colored in red. **(F)** Pathway enrichment (Enrichr) on tRCC-selective gene dependencies (relative to ccRCC) as experimentally determined by CRISPR screening of three tRCC lines (Fig. 4E). **(G)** Protein-protein interaction network (STRINGdb) amongst OXPHOS-related tRCC-selective gene dependencies (vs. ccRCC).

**Fig. 5.**
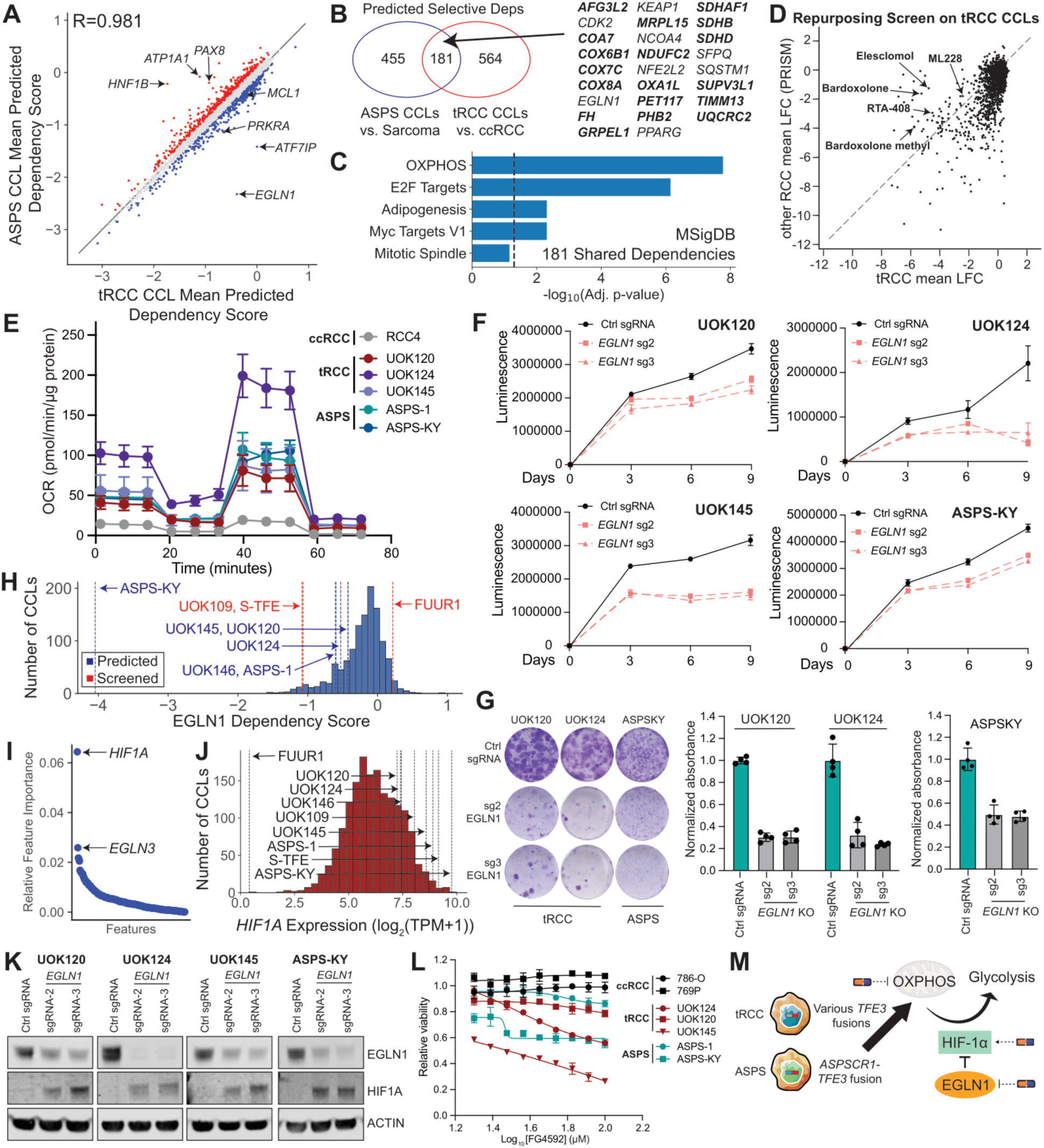
Virtual CRISPR screening nominates *EGLN1* as a shared dependency across TFE3 fusion driven tumors. **(A)** Predicted dependency scores for 6,283 PD genes in ASPS cell lines (y-axis; mean of 2 cell lines) vs. tRCC cell lines (x-axis; mean of 7 cell lines). Dependencies predicted to be tRCC-selective are labeled in red while ASPS-selective dependencies are in blue. **(B)** Venn diagram comparing tRCC-selective (relative to ccRCC) and ASPS-selective (relative to other sarcomas) predicted dependencies. Bolded genes are involved in mitochondrial metabolism/oxidative phosphorylation. *P*=1.65e-36 by hypergeometric test. **(C)** Pathway enrichment (Enrichr) on all 181 shared predicted ASPS/tRCC-selective dependencies from Fig. 5B. **(D)** Mean log_2_(fold-changes) in viability following compound treatment (2.5 μM, 5 days) in four tRCC cell lines (FUUR-1, S-TFE, UOK109, UOK146; x-axis) screened with the PRISM drug repurposing library compared to six previously screened ccRCC cell lines with same library in DepMap (y-axis). **(E)** Seahorse metabolic flux analysis in tRCC and ASPS cell lines. The ccRCC cell line RCC4 serves as a highly glycolytic control. **(F)** Cell viability after *EGLN1* knockout across a panel of tRCC (UOK120, UOK124, UOK145) and ASPS (ASPS-KY) cell lines. Shown as mean +/- s.d., n=3 biological replicates per condition. **(G)** Representative image (left) and quantification of colony formation assay (right) for tRCC (UOK120, UOK124) or ASPS cells (ASPS-KY) after *EGLN1* knockout. Shown as mean +/- s.d., n=3 biological replicates per condition. **(H)** Distribution of *EGLN1* Chronos score across all cell lines in DepMap (N=1207). Predicted dependency scores (blue) or screened Chronos scores (red) for tRCC/ASPS cell lines are shown as dotted lines. **(I)** Ranked relative feature importance for RNA predictors of *EGLN1* dependency score. **(J)** Histogram of *HIF1A* expression across all cell lines in DepMap (N=1207); expression in tRCC/ASPS cell lines are indicated as dashed lines. **(K)** Western blot showing stabilization of HIF1α protein after *EGLN1* knockout in tRCC/ASPS lines. **(L)** ccRCC (786O, 769P), tRCC (UOK124, UOK120, UOK145), or ASPS cell lines (ASPS-1, ASPS-KY) treated with indicated concentrations of pan-EGLN inhibitor FG-4592 and assayed for cell viability after 3 days with CellTiter-Glo. Viability at each concentration is relative to vehicle-treated cells, shown as mean +/- s.d., n=3 biological replicates. **(M)** Model schematizing how TFE3 fusion-driven OXPHOS sensitizes two distinct cancers – tRCC and ASPS – to EGLN inhibition/HIF-1α activation.

To integrate these predictions with unbiased functional screens, we then performed genome-scale pooled CRISPR-Cas9 knockout screening (Broad Brunello library (*156*)) in three tRCC cell lines representing two distinct *TFE3* fusions (FUUR-1: *ASPSCR1-TFE3*; S-TFE: *ASPSCR1-TFE3*; UOK109: *NONO-TFE3*) (**Fig. S6D**). To calibrate against DepMap, we calculated Chronos dependency scores from our screen (**Table S3**, see **Methods**) and found excellent concordance between our experimentally-derived Chronos scores and predicted dependency scores in all tRCC cell lines screened, demonstrating that our method can accurately predict the dependency landscape of unscreened samples from RNA-seq, even types not present in the original training data (**Fig. 4D**).

We then compared experimentally-derived Chronos scores for each gene in tRCC cells (averaged across the 3 tRCC cell lines screened) to Chronos scores for these same genes in either clear-cell renal cell carcinoma (ccRCC) cell lines or all cancer lineages screened in the DepMap (**Fig. 4E, Fig. S6E**). Among the most selective dependencies in tRCC cells were *TFE3* and *ASPSCR1* (fusion partner of *TFE3* in two of three cell lines screened), although neither of these genes were PD genes likely due to the complete absence of *TFE3* fusion cell lines in the DepMap training data (**Fig. 4E**). Selective essentiality of the *TFE3* fusion in tRCC was validated across a panel of cell lines in a growth competition assay (**Fig. S6F-H**) (*24*, *157*). Together, these results indicate that the driver *TFE3* fusion represents the primary selective essentiality in tRCC and that wild type *TFE3* is dispensable in non-fusion cancer cells.

Kidney lineage-defining transcription factors (*PAX8*, *HNF1B*) (*158–160*) were strong dependencies in both tRCC and ccRCC and dependency correlated strongly with their expression (**Fig. S6B-C, Fig. S7A**). Accordingly, both genes were highly expressed in tRCC and ccRCC but not in other *TFE3*-driven malignancies (melanotic kidney tumors, PEComa, ASPS) that may be of mesenchymal origin (**Fig. S7B-F**). Three genes involved in mevalonate synthesis (*PMVK, MVK, MVD*) were also strong dependencies in tRCC and had variable levels of dependency across ccRCC cell lines (**Fig. S7G**); given prior reports of perturbed cholesterol biosynthesis in ccRCC, this may also represent a form of pan-RCC lineage dependency that holds across various types of kidney cancer (*161*, *162*).

Finally, we performed gene ontology enrichment (*163*) on dependencies selectively essential to tRCC vs. ccRCC cell lines based on our functional screens. The enriched pathways remarkably mirrored our dependency predictions on tRCC tumors (**Fig. 4C**, **Fig. 4F, Fig. S8A-B**), with pathways related to oxidative phosphorylation, mitochondrial translation, and electron transport strongly enriched amongst tRCC-selective dependencies (**Fig. 4E-G**). Specific dependencies included: (**1**) genes involved in the transcription and translation of mitochondrially-encoded genes (*POLRMT*: mitochondrial RNA Polymerase that transcribes mitochondrial DNA (mtDNA); *MRPL48*: component of the mitochondrial ribosome (“mitoribosome”); *ERAL1*: involved in mitochondrial rRNA assembly; *NARS2*: mitochondrial asparaginyl tRNA synthetase); (**2**) genes encoding enzymes in the citric acid (TCA) cycle (*SDHA, SDHB*); (**3**) genes encoding components of the mitochondrial ATP synthase and electron transport chain (*ATP5F1A, ATP5F1D, ATP5F1E, ATP5ME, ATP5PD, CYCS*); (**4**) genes involved in the assembly or biogenesis of iron-sulfur clusters, which are critical for Complex I, II and III activity within the electron transport chain (*164*) (*FDX2, HSCB, ISCA1*) (**Fig. 4E-G**, **Fig. S8A**). We functionally validated that knockout of several of these genes (*ISCA1, SDHA*, *MRPL48, POLRMT*) selectively impairs the growth of tRCC cells in assays for cell proliferation, cell viability, and clonogenic capacity (**Fig. S8C-H**).

Altogether, these results indicate that our dependency predictions on tRCC tumor and cell line transcriptomes and unbiased functional screens converge on a core group of tRCC-selective vulnerabilities related to oxidative phosphorylation.

### Dependency prediction converges with small molecule and genetic screens to implicate the EGLN-HIF-1α axis as a vulnerability in two distinct TFE3 fusion-driven cancers

Given the excellent alignment of our dependency prediction and unbiased functional approaches in tRCC, we next sought to interrogate other *TFE3* fusion-driven cancers. *TFE3* fusions drive a spectrum of rare cancers apart from tRCC, including alveolar soft part sarcoma (ASPS), some endothelial hemangioendotheliomas (EHE), and some perivascular epithelioid cell tumors (PEComa) (*165–167*). Most of these diseases have been scarcely profiled and have limited or no models amenable to high-throughput screening. While all of these cancers, along with tRCC, share *TFE3* driver fusions, they may differ in terms of the cell of origin as well as in co-occurring genetic driver alterations, which may modulate dependency profile.

We sought to compare the dependency profile of tRCC with ASPS, which accounts for <100 new cases annually in the United States, has few robust models, and has been scarcely profiled (*168*). We performed dependency predictions on RNA-Seq data from seven ASPS tumors (*126*) and two ASPS cell lines (ASPS-1, ASPS-KY). Although the dependency profiles of tRCC and ASPS cell lines were largely concordant (R=0.981), several dependencies were predicted to be selective for ASPS cells versus tRCC, including *MCL1* and *PRKRA* (**Fig. 5A-C**), which we validated genetically (via CRISPR-mediated knockout of *MCL1* and *PRKRA*) and chemically (using the clinical-grade MCL1 inhibitor murizatoclax (*169*)) in a panel of tRCC and ASPS cell lines. (**Fig. S9A-F**). Our model identified low *BCL2L1* (which encodes the anti-apoptotic factor BCL-xL) expression as a top feature predicting of *MCL1* sensitivity (**Fig. S9G-H)**; accordingly, *BCL2L1* expression was lower in ASPS cell lines/tumors versus tRCC cell lines/tumors and kidney cancer-adjacent normal tissue (**Fig. S9I)**. Conversely, and consistent with tRCCs being of renal epithelial origin and ASPS being of a mesenchymal origin, the renal lineage factors *PAX8* and *HNF1B* were not predicted as dependencies in ASPS (**Fig. 5A**, **Fig. S9A**). We conclude that tRCC and ASPS harbor some distinct selective vulnerabilities, despite both sharing the same driver fusion.

We next investigated shared dependencies between tRCC and ASPS. We identified selective dependencies in tRCC (relative to ccRCC) and ASPS (relative to other sarcomas) and observed significant overlap in these genes (*P*=1.65e-36, **Fig. 5B**). Shared dependencies between tRCC and ASPS were enriched for genes involved in oxidative phosphorylation and mitochondrial metabolism, consistent with prior transcriptional profiling of ASPS (*170*) (**Fig. 5C**). In addition, we observed shared dependency on the prolyl hydroxylase *EGLN1* (predicted to be quantitatively stronger in ASPS than tRCC) (**Fig. 5A-B**). This finding is consistent with our recent model that OXPHOS-dependent tRCCs are sensitive to inhibition of *EGLN1*, which stabilizes its substrate HIF-1α and induces metabolic reprogramming away from OXPHOS and towards glycolysis (*155*, *171*).

As multiple processes involved in mitochondrial metabolism, redox, and OXPHOS can be perturbed pharmacologically, we next sought to determine whether an unbiased small molecule screen would recapitulate our predicted genetic dependencies. We performed a viability screen of a Drug Repurposing Library (*172*) (∼7000 compounds, encompassing FDA/globally approved compounds, clinical stage compounds, and preclinical compounds at a single 2.5 μM dose) in four tRCC cell lines (FUUR-1, S-TFE, UOK109, UOK146; ASPS cell lines omitted due to difficulty in culturing for high-throughput assays). We compared results to six RCC cell lines screened with the same chemical library on the PRISM platform (*66*). Strikingly, several compounds scoring as selective for tRCC aligned with our functional genetic data in tRCC and our dependency predictions for both tRCC and ASPS. These included: (**1**) the NRF2 activators bardoxolone, bardoxolone methyl, and RTA-408 (which would be predicted to phenocopy *KEAP1* knockout, predicted tRCC selective vs. ccRCC [**Fig. S4D, Fig. 5B**], and induce reductive stress, a vulnerability in OXPHOS-high cells) (*155*, *173*); (**2**) the mitochondrial poison elesclomol, to which mitochondrially-primed cells are uniquely sensitive (*174*, *175*); (**3**) the HIF pathway activator ML228 (*176*), which would be predicted to phenocopy *EGLN1* inhibition and suppress OXPHOS by inducing glycolytic reprogramming, as described above (**Fig. 5D**).

To validate these multiple orthogonal lines of data linking OXPHOS vulnerability to two TFE3 fusion-dependent contexts, we first performed Seahorse metabolic flux analysis and confirmed that ASPS cells exhibit an increased oxygen consumption rate (OCR) relative to ccRCC, consistent with what we have recently reported for tRCC (*155*) and in line with their presumably increased mitochondrial capacity (**Fig. 5E**). Genetic knockout of *EGLN1* uniformly impaired cell proliferation in multiple tRCC and ASPS cell lines tested despite there being only 90/1207 dependent lines (7.5% with *EGLN1* Chronos score ≤ -0.75) across the entire DepMap (**Fig. 5F-H**). *HIF1A* expression represents the top predictive feature of *EGLN1* dependency in our model, presumably because high *HIF1A*-expressing, high OXPHOS cancers would be most susceptible to HIF-1α protein accumulation upon EGLN1 inhibition; the second predictive feature is *EGLN3* expression, presumably due to redundancy between *EGLN1* and *EGLN3* (*177*). Both tRCC and ASPS cell lines express *HIF1A* at high levels compared to other lineages in DepMap (with the exception of FU-UR-1), with ASPS-KY displaying the second highest expression of all cell lines (**Fig. 5I-J**). Inhibition of *EGLN1* was accompanied by stabilization of HIF-1α in ASPS, which promotes glycolytic reprograming and would be expected to be deleterious to OXPHOS-dependent cells (*178*) (**Fig. 5K**). These effects were also recapitulated with the pharmacologic EGLN inhibitor, FG-4592, in both tRCC and ASPS cells (**Fig. 5L**).

Overall, these findings – which we identified via dependency prediction and subsequently validated experimentally – are consistent with TFE3 fusions rewiring the metabolic program of both tRCC and ASPS to oxidative phosphorylation, rendering these tumor types uniquely dependent on mitochondrial metabolism and therefore sensitive to pharmacologic HIF-1α activation (**Fig. 5M**).

### An interactive portal for exploring candidate dependencies

To facilitate the exploration of our dependency prediction data, we have established a web portal to explore TrPLet dependency predictions (http://viswanathanlab-deps.dana-farber.org/). Through this website, users can: (**1**) review predicted dependency scores for PD genes across the TCGA and multiple other tumor and cell line RNA-Seq cohorts, in comparison to experimentally-derived dependency scores from the DepMap (**2**) view the comparative performance of different PD gene prediction models (**3**) identify expression features associated with the prediction of dependency score for each PD gene. Instructions to run the TrPLet pipeline for dependency prediction on user-supplied RNA-Seq data are also provided.

## Discussion

Recent functional genetic studies have proven an effective engine for target discovery, but many rare cancers and even some common cancer models have been underrepresented in these efforts. We attempted to bridge this gap by pursuing the alternative approach of predicting a tumor’s dependency landscape via its transcriptional profile. Prior studies have reported approaches to predict a tumor’s dependency profile or infer synthetic lethality by virtue of its transcriptional and/or genomic features (*47–49*, *51*) but each differs somewhat in methodology and performance and these models have not typically been extensively validated on unseen cancer types. Our approach, which predicts dependencies on substantially more genes than prior models, utilizes expression features that can be readily obtained by clinical transcriptome sequencing of tumor tissue. We suggest that this type of approach can be broadly employed for precision medicine applications across tumor types, and to guide treatment selection in rare cancers, for which there may be no evidence-based standard of care.

Our dependency predictions are robust to functional validation across multiple contexts, including selective dependencies for individual tumor types not profiled in the training set (e.g. for tRCC, FH-RCC, and ASPS), synthetic lethal relationships (e.g. associated with *RB1* inactivation or *PPP2CA* loss), and predicting response to targeted therapies (e.g. KRAS G12C inhibitors). Importantly, these relationships, and others previously identified via functional screens (e.g. synthetic lethal targets with MSI-H), were inferred directly from RNA-sequencing of tumor samples. Moreover, several of these (e.g. *RB1* synthetic essentials) reach significance only via a “virtual” CRISPR screening approach and not when analyzing experimentally screened samples, presumably due to increased power from larger tumor transcriptome datasets. These findings are consistent with prior reports that synthetic essential relationships may be evinced by transcriptional features (*51*, *92*) and highlight the tractability of transcriptome-driven dependency discovery.

A prime application of our approach is in defining dependencies in rare or understudied cancer types. For example, although almost all discovery biology in kidney cancer has to date has focused on ccRCC, drugs for ccRCC are typically less effective in non-ccRCCs (*179*), which are driven by distinct biology. By applying dependency prediction to a spectrum of kidney cancers – most of which have not been systematically characterized – we suggest that the several dozen histologic types of kidney cancer (*17*) collapse into five distinct groups in dependency space. Dependencies related to energy metabolism encapsulate this notion: while deficiency in TCA cycle enzymes such as FH and SDHA/B drives tumorigenesis in glycolytic renal cancers (e.g. FH-RCC and ccRCC), these same genes represent dependencies in high OXPHOS renal cancers such as tRCC (*180*). We nominate several potentially actionable dependencies in different RCC subtypes including *NFE2L2* in FH-RCC, *USP1* in oncocytic tumors, and β-catenin signaling in Wilms’ tumor. Our findings underscore the importance of molecularly-informed, subtype-specific treatments in kidney cancer.

Our approach provides a scalable solution to therapeutic targeting in rare cancers with limited cell line models, which we exemplified using the case of *TFE3* fusion cancers. In the case of tRCC, over fifteen different *TFE3* fusion partners have been reported (*24*), but existing cell line models represent only about five. For ASPS, a much rarer cancer, only two cell lines exist but both are slow growing and challenging to culture at the scale required for genome-scale screening. Other *TFE3* fusion-driven cancers are even rarer, without existing models at all. Here, by an integrative approach that included dependency prediction on tumor samples, unbiased functional genetic and small molecule screening in amenable models, and targeted functional validation in more challenging ones, we identified and extensively validated the OXPHOS pathway as a shared dependency of two distinct *TFE3* fusion cancers: tRCC and ASPS. Our genome-scale functional screens in tRCC pinpoint multiple targetable entry points involved in mitochondrial metabolism and oxidative phosphorylation, including components of the citric acid (TCA) cycle, mitochondrial transcription and translation, and the electron transport chain; these screens also dovetail remarkably with two recent studies demonstrating that TFE3 fusions metabolically rewire tRCCs towards oxidative phosphorylation (OXPHOS) in tRCC (*155*, *171*) and are consistent with the role of wild type TFE proteins as critical regulators of energy metabolism (*181–183*). We also point to HIF-1α activation, which can be achieved pharmacologically, as a therapeutically tractable strategy to enable OXPHOS inhibition in these two cancers of unmet need.

Overall, we suggest that our combined approach of functional screening and dependency prediction may catalyze precision oncology in many settings – including those where experimental models may be limited or where discovery biology efforts are resource-limited – by enabling the inference of genetic dependency, synthetic lethal relationships, and drug responses directly from patient tumors.

## Acknowledgements

Schematic illustrations were created with BioRender.com.

## Author contributions

S.R.V designed and supervised the study. A.S., B.L., J.L., S.R.V wrote the manuscript with input from all co-authors. A.S. performed analysis of genome-scale CRISPR screening data, dependency prediction and developed the predictive model used in this study. B.L. and J.L. led experimental work including genome-scale CRISPR screening and dependency validation. R.D., D.Y., Y.W, D.R.T., M.T., C.N.W. contributed to experimental work related to dependency validation. Y.C. and P.K. assisted in analysis of dependency data and establishment of the web portal. C.B. and J.H.C performed or supervised small molecule screen. T.K.C. provided input regarding kidney cancer subtyping. J.G.D. contributed to design of screening and validation studies. B.J.D. provided input on the overall study design and supervised validation studies related to Rb synthetic essentiality.

## Data and materials availability

Chronos scores from tRCC CRISPR screen are available in **Table S3.** External datasets analyzed are public and are available from the respective cited publications. Predicted dependency scores for all external datasets are available in **Table S2**, which is deposited in Dropbox due to its large size: https://www.dropbox.com/scl/fi/8o3d0yqttmjzjh6i84vzq/Table_S2.xlsx?rlkey=qlthlp1er8uiirs7yjogolwpl&st=0epypnmb&dl=0. Interactive visualization of dependency data is available in the TrPLet portal: http://viswanathanlab-deps.dana-farber.org/. Code for the developed tool is publicly available in Github: https://github.com/SViswanathanLab/TrPLet.

## Materials and Methods

### Dependency Prediction Models

#### Model development and evaluation

The DepMap 23Q2 expression matrix (converted to log_2_(TPM+1), https://depmap.org) and 23Q2 CRISPR-KO dependency score matrix (Chronos-normalized, https://depmap.org) were downloaded and subset to shared cell lines. We split these data into 5 equal subsets for 5-fold cross-validation (four training folds and evaluation on a validation fold). Expression was Z-scored for each gene. For each gene, 60 models were trained to predict its dependency score (5 model types, 12 levels of dimensionality reduction). The 5 model types tested were elastic net regression, ridge regression, lasso regression, SVR with a linear kernel, and SVR with an RBF kernel. To reduce dimensionality prior to training/testing, we calculated the Pearson correlation coefficient between each feature (i.e. Z-scored gene expression) and the dependency score of the gene being analyzed in the training data and restricted to the top N features (with highest absolute value of Pearson correlation coefficient). For each model type, we tested the following values for N: 1, 2, 3, 5, 10, 50, 100, 500, 1000, 2500, 5000, 10000 (**Figure S1**). We selected the best performing model for each gene dependency being predicted (highest cross-validated performance).

Performance was assessed through correlation coefficient between predicted dependency scores and CRISPR screen-derived Chronos scores, averaged across all the validation folds. We utilized the previously established performance threshold of R ≥ 0.2 to define predictable genes (*47*). RNA-only dependency prediction models were trained for 16,845 genes, using the best model for each gene. Of these, 6,283 genes were predictable at the performance threshold of R ≥ 0.2 (“PD genes”) (**Fig. 1C, Fig. S1**). This significantly outperformed prior studies, which did not optimize model type for each gene, resulting in >3x as many PD genes as the prior state-of-the-art (N=1,966 at R ≥ 0.2 (*47*) **Fig. S1**). Furthermore, across 1,298 DepOIs reported in Chiu et al. we observed superior performance (mean R=0.37 for TrPLet vs. mean R=0.18 for Chiu et al.) (**Fig. S1**) (*48*).

The average Pearson correlation coefficient for predicted dependency scores vs. screened Chronos scores was R=0.20 across 16,845 genes. The correlation between predicted dependency scores vs. screened Chronos scores across all gene-cancer cell line pairs in the test data was R=0.92. Specifics and code for this project, a snakemake pipeline that predicts dependencies from RNA-seq fastq files or count matrices, as well as scripts for interactive visualization of predicted dependencies in this manuscript are available in Github: https://github.com/SViswanathanLab/TrPLet. An interactive visualization of all predicted dependencies in this paper is available in a web portal: http://viswanathanlab-deps.dana-farber.org/. The best model type for each gene is available in **Table S1**. For models in **Fig. 1B** including mutation calls, we utilized binarized hotspot mutation calls (https://depmap.org) that were merged with expression features prior to training/testing.

#### Model deployment

The models for each gene were retrained on the entire DepMap dataset prior to testing on external datasets. We extracted model coefficients directly from sklearn when possible. To calculate approximate coefficients for SVR RBF, we used a kernel trick taking the linear combination of support vector weighted by dual coefficients from the RBF kernel. Broadly, our model was applied to three types of data: (1) TCGA tumor RNA-seq, (2) non-TCGA tumor RNA-seq, (3) cell line RNA-seq. (1) For TCGA tumor RNA-seq, we Z-scored the expression of each gene (log_2_(TPM+1)) within TCGA and predicted on the resulting normalized expression data. Clustering (two-component t-SNE) based on dependency scores (predicted: TCGA, experimentally-derived via CRISPR screen: DepMap) using this approach resulted in TCGA tumors clustering with screened cell lines from DepMap of the same lineage (see **Fig. 1D**). (2) For non-TCGA tumor RNA-seq, we downloaded RNA-seq fastq files or count matrices when present, from the Gene Expression Omnibus (GEO). Reads were aligned to GENCODE v38 transcript reference using STAR/RSEM (*185*, *186*). The resulting count matrices were inner joined with the TCGA count matrix (*187*) (https://osf.io/gqrz9/files/osfstorage), and batch corrected using ComBat-seq (*188*) using lineage as a covariate (for this purpose, tRCCs, Wilms’ tumors, CDC, and RMC were classified as “KIRP”; ASPS, PEComa, and EHE were classified as “SARC”; ccRCC was classified as “KIRC”). The counts were then normalized to gene-level transcripts per kilobase million (TPM), converted to log_2_(TPM+1), and each gene’s expression was then Z-scored (across the combined matrix consisting of the external dataset and TCGA). The resulting Z-scored expression in the external dataset was then used for prediction, as described above. (3) For cell line RNA-seq, a gene-level normalized expression matrix (log_2_(TPM+1)) was either downloaded or generated from STAR/RSEM. The expression of each gene in the resulting matrix was scaled (Z-scored) using the mean/standard deviation calculated when scaling the DepMap expression matrix. The resulting Z-scored expression values were used for prediction. Batch correction was forgone in this use-case based on tSNE clustering of tRCC cell lines with kidney cell lines in DepMap, and ASPS cell lines with soft tissue and CNS cell lines in DepMap based on expression profile. Unless otherwise specified, a predicted selective dependency has threshold |Δ(mean predicted dependency score)| > 0.1 between the groups being compared.

### Cell Lines

H460 (ATCC® HTB-177), PC3 (ATCC® CRL-1435), 786-O (ATCC® CRL-1932™), 293T (ATCC® CRL-11268™), Caki-1 (ATCC® HTB-46™), HT29 (ATCC® HTB-38), HCC38 (ATCC® CRL-2314), NCI-H358 (ATCC® CRL-5807), A549 (ATCC® CRM-CCL-185), NCI-H1048 (ATCC® CRL-5853), HCC827 (ATCC® CRL-2868), NCI-H1975 (ATCC® CRL-5908), CORL311 (Sigma Aldrich, 96020721), NCI-H526 (ATCC® CRL-5811), CORL47 (Sigma Aldrich, 92031915), NCI-H838 (Hamon Cancer Center), NCI-H2009 (Hamon Cancer Center), NCI-H2228 (Hamon Cancer Center), UOK109 (Dr. W. Marston Linehan’s laboratory, National Cancer Institute), UOK146 (Dr. W. Marston Linehan’s laboratory, National Cancer Institute), s-TFE (RIKEN, # RCB4699), UOK120 (Dr. W. Marston Linehan’s laboratory, National Cancer Institute), UOK124 (Dr. W. Marston Linehan’s laboratory, National Cancer Institute), UOK145 (Dr. W. Marston Linehan’s laboratory, National Cancer Institute), UOK262 (Dr. W. Marston Linehan’s laboratory, National Cancer Institute), ASPS1 (National Cancer Institute (*189*)), and ASPS-KY (Riken, RCB5683) cell lines were cultured at 37°C in DMEM with 10% FBS, 100 U/mL penicillin, and 100 μg/mL Normocin (Thermo Fisher: #NC9390718). HCC827, NCIH1975, NCIH358, NCIH838, NCIH2009, and NCIH2228 were also cultured in RPMI with 5% FBS. The FU-UR-1 (Dr. Masako Ishiguro’s laboratory, Fukuoka University School of Medicine) cell line was cultured at 37°C in DMEM/F12 (1:1) with 10% FBS, 100 U/mL penicillin, and 100 μg/mL Normocin.

### Genome-Scale CRISPR Knockout Screens

For the UOK109, FU-UR-1, and s-TFE cell lines, Cas9-expressing cells were constructed as follows: each parental cell line was seeded in 12-well plates (2.5×10^5^ cells/well) and incubated at 37°C overnight. The following day, the medium was replaced, and cells were incubated with lentivirus corresponding to the pLX_311-Cas9 plasmid (Addgene plasmid #96924), which encodes the Cas9 protein, and 0.8 μg/mL polybrene. After overnight incubation at 37°C, the cells were trypsinized the following day and cultured in selective media containing 5 μg/mL blasticidin. After selection, Cas9 expression and activity were confirmed in each transduced cell line via western blotting and a Cas9-activity assay as described in a previous reference (*190*).

The Broad Institute Brunello sgRNA library (77,441 sgRNAs targeting 19,114 genes with 1,000 non-targeting control sgRNAs) was applied for the CRISPR Screen (*191*, *192*). UOK109, FU-UR-1, and s-TFE cells were seeded into 12-well plates at a density of 1.5/1.25/1.5×10^6^ cells/well, with 1.2 µg/mL polybrene and virus titrated at MOI <0.3 and spun at 1000g for 2 hours at 33°C. After spinfection, 1 mL medium was added to each well and incubated at 37°C overnight. The following day, all cells were trypsinized and expanded into 15 cm plates at 4×10^6^ cells/plate with 4/5/5 µg/mL puromycin for a week. Medium with puromycin was replaced every 3 days. After puromycin selection, cells were seeded at 3×10^6^ cells/plate and replated every 7 days in 15 cm plates for 21 days. At 28 days post-infection, cells were collected and stored at -20°C before genomic DNA was collected.

Genomic DNA was collected with Takara NucleoSpin Blood Kits (Macherey-Nagel), following the manufacturer’s protocol. Before sequencing, genomic DNA samples were amplified by PCR.

### Chronos Score Calculation

We calibrated our results against the DepMap (*13*, *36*) by calculating a Chronos score for each gene assayed in our screens. This metric represents the relative essentiality of a gene accounting for various potential confounders, including sgRNA efficiency, copy number related bias, and heterogenous cutting events (**Methods**) (*37*); by the Chronos metric, cell-essential genes have a score of approximately -1 while non-essential genes have a score of approximately 0 (*37*). Essential genes (*184*) had a mean Chronos score of –0.930 across tRCC cell lines screened in our study while non-essential genes had a mean Chronos score of -0.026, suggesting that informative comparisons can be made between our data and those generated via the DepMap despite some differences in protocol and library (**Methods**). Importantly, while cell lines in the DepMap effort were screened using the Avana CRISPR knockout sgRNA library (*156*) and the tRCC cell lines in this study utilized the Brunello library, the strong concordance in the scores for both essential and non-essential genes suggests that informative comparisons can be made between our data and those generated via the DepMap effort, despite the use of different genome-scale sgRNA libraries and the screens being conducted over different durations.

To calculate Chronos scores, log_2_(fold-changes) in sgRNA abundance on day 28 of the screen were calculated using MAGeCK (*193*), using plasmid DNA as a reference. Exome sequencing data was aligned to hg38 using bwa mem (*194*), and copy-number was calculated using PureCN (*195*), as previously described (*157*). Chronos was used to normalize log_2_(fold-changes) in sgRNAs with segmental copy-number correction (*37*). ccRCC cell lines were defined based on Cellosaurus NCIt disease type and included: OSRC2, CAKI2, SLR23, KMRC20, UOK101, SLR24, KMRC3, CAKI1, TUHR10TKB, SLR26, KMRC1, KMRC2, SNU349, UMRC3, and RCCFG2.

### Lentiviral Production

All sgRNAs were cloned into plentiCRISPRv2 (Addgene, #52961, puromycin resistance) as described (*196*, *197*). For RB1 colony formation assays and isogenic drug assays, pLV[CRISPR]-hCas9:T2A:Hygro-U6 was used as a backbone. For NFE2L2 knockdown, dox-inducible shRNA was cloned into a Gateway-compatible lentiviral vector as previously described (*198*). sgRNA/shRNA sequences are listed in **Supplementary Table 4**. All the constructs were confirmed by Sanger/whole plasmid sequencing.

Lentivirus was prepared by transfecting 293T cells with three plasmids (plentiCRISPRv2, psPAX2, and pMD2.G) using polyethylenimine (PEI). Media was replaced with standard growth media after 12 hours, and supernatant containing the virus was collected 48 hours post-transfection.

### Validation of Genome-Scale CRISPR-Cas9 Screens and Dependency Predictions

Cell lines were transduced with lentivirus expressing CRISPR-Cas9 and sgRNA targeting the gene of interest, selected by puromycin/hygromycin B, and then seeded in 96-well plates for confluence and proliferation assays with cell densities of 400-2,000 cells/well depending on the cell line. On days 3-28, depending on the cell line, cell growth medium was removed from the plates and the Cell Titer Glo reagent (Promega, #G7571) was added following the user’s instructions. Plates were then shaken at room temperature for 10 minutes. Luminescence was measured on a SpectraMax plate reader. For drug assays, cells were incubated with milademetan (MCE: #HY-101266), murizatoclax (MCE: #HY-109184), SKPin C1 (MCE: #HY-16661), or FG-4592 (MCE: # HY-13426) for the duration of time indicated. For cell confluence assays, confluence on each plate was determined using a Celigo Imaging Cytometer daily.

### Competition Assay

Caki-1, 786-O, UOK109, FU-UR-1, and s-TFE were transduced with lentivirus expressing EGFP, CRISPR-Cas9, and sgRNA (either control sgRNA (098) or sgRNA against TFE3). After 3 days, the EGFP-positive rate was measured by a Fortessa flow cytometer to ensure it was higher than 90%. Seven days after viral infection, viral-infected cells were mixed with non-infected parental cells in a 1:1 ratio. Mixed cells were plated in 6-well plates. On days 3-24, cells were trypsinized and resuspended in 4% FACS buffer (FBS/PBS), and the EGFP-positive rate was measured by a Fortessa flow cytometer. All flow cytometry data were analyzed with FlowJo. EGFP positive percentage in each condition at each time point was first normalized to value in that condition at d0, and then normalized to sgControl.

### Colony Formation Assays

Cell lines were transduced with lentivirus expressing CRISPR-Cas9 and sgRNA targeting the gene of interest, selected by puromycin, and then seeded in 12-well plates at various cell densities of 500-6,000 cells/well depending on the cell line. Media was replaced every 7 days. After 10-28 days, medium was aspirated, and cells were fixed and stained with 0.5% crystal violet in 25% (volume) methanol solution for about 15 minutes. Stained cells were washed with water and air-dried. Plates were scanned with an Epson scanner and quantified using ImageJ.

### Immunoblotting

Cells were lysed in RIPA buffer (Thermo Fisher Scientific, #89901) supplemented with protease and phosphatase inhibitors (Roche, #11836170001, Roche, #4906845001). Protein concentrations were determined using the BCA assay (Thermo Fisher Scientific, #23225). Equal amounts of protein were resolved by NuPAGE 4-12% Bis-Tris Protein Gels (Thermo Fisher Scientific, #NP0335), Proteins were transferred to nitrocellulose membranes (Life Technologies, #IB23001) using an iBlot2 (Thermo Fisher Scientific). Membranes were blocked with blocking buffer (Thermo Fisher Scientific, # NC1660550). Immunoblot analysis was performed with the indicated primary antibodies in antibody dilution buffer (Thermo Fisher Scientific, #NC1703226) overnight at 4°C. Membranes were incubated with LI-COR secondary antibodies in antibody dilution buffer. Protein bands were visualized using the Odyssey Clx Infrared Imaging System (LI-COR Biosciences). Specifically, Western blot for sample collection of HIF-1α, cells were seeded in equal numbers in 6-well plate on Day 1. On Day 2, cells were harvested in equal volumes of Laemmli buffer, followed by boiling at 95°C for 10 minutes. Primary antibodies used were: Nrf2 (Abcam, ab62352), RB1 (CST, #9309), β-actin (CST, #3700), GAPDH (CST, #97166), HIF-1α (Fisher Scientific, BDB610959). <u>Drug Screen</u>

TRCC cell lines (FU-UR-1, UOK-109, UOK-146, S-TFE) were screened against the Broad Repurposing Library (https://repo-hub.broadinstitute.org/repurposing) (*172*). This library of ∼6800 compounds is a curated collection of FDA-approved drugs, clinical trial drugs, and pre-clinical tool compounds, which are deeply annotated for phase, target, indication and mechanism-of-action. Assay-ready plates of this library were created by printing nanoliter quantities of each drug into 384-well Corning 3570BC plates using a Labcyte Echo acoustic liquid handler.

The lines were grown in DMEM (Gibco #11965-084) + 10% Fetal Bovine Serum (Sigma #F2442-500ML) + 1X sodium pyruvate (ThermoFisher #11360-070) + 1X PenStrep (ThermoFisher #15140-163). 50 µL of cells at the optimized cell density per cell line (FU-UR-1: 500 cells/well, S-TFE: 1000 cells/well, UOK-109: 1000 cells/well, UOK-146: 750 cells/well) was added to the assay-ready plates using a Combi multidrop dispenser and the compounds were screened in duplicate at a final assay concentration of 2.5 µM for 72 hours. Cell Titer-Glo was reconstituted per the manufacturer’s protocol and added to each well (10 µL) using a Combi multidrop dispenser. Cell Titer-Glo uses the ATP levels in the cell as a surrogate for viability, driving a luminescence reaction. After a 10 minute incubation, the luminescence signal was readout on a BMG PheraStar plate reader.

The raw luminescence data was analyzed using Genedata. Each compound effect was normalized to DMSO and reported as a % activity; this was converted to log_2_(fold-change): LFC = log_2_(1-X) for comparison against DepMap ccRCC cell lines profiled by the library (786O, OSRC2, RCC10RGB, KMRC3, TUHR4TKB, KMRC1).

### Seahorse Assay

Seahorse assay was performed as previously described (*155*). Oxygen consumption rates (OCR) were determined with the XF Cell Mito Stress Kit (Agilent: #103015-100). ccRCC cell line (RCC4), tRCC cell lines (UOK120, UOK124 and UOK145) and ASPS cell lines ( ASPS-1 and ASPS-KY) were seeded on a poly-L-lysine coated 96-well Seahorse plate (Agilent: #101085-004) a. Cells were incubated overnight at 37 ℃ in 5% CO2 incubator. XF96 FluxPak sensor cartridge was hydrated according to manufacturer’s instructions. During the following days, the growth medium was removed, and cells were washed with pre-warmed seahorse medium (XF DMEM (Agilent: #103575-100) supplemented with 10 mM glucose, 1 mM pyruvate solution, and 2 mM glutamine). After washing, cells were incubated in 180 μL Seahorse medium at 37°C in non-CO2 incubator for 45-60 min. The oxygen consumption rates were measured by XFe 96 extracellular flux analyzer by adding oligomycin, FCCP and rotenone/antimycin A to each cartridge port. All OCR values were normalized to total protein content as measured by BCA (Thermo Fisher Scientific, #23225) according to manufacturer’s instructions.

## Supplementary Figures

**Fig. S1.**
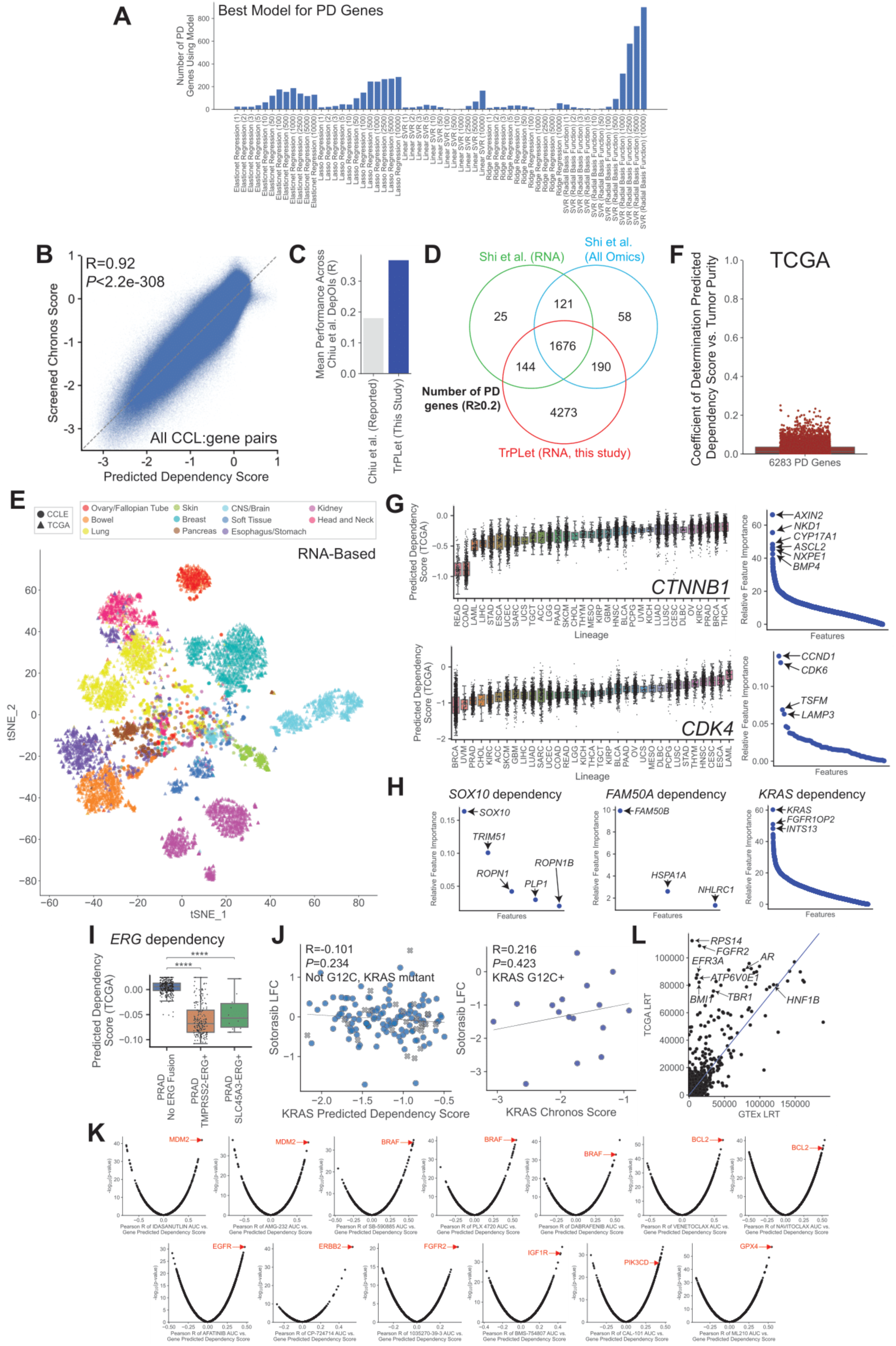
Performance metrics for dependency prediction and application to RNA-seq datasets. **(A)** Best performing model type for each of the 6,283 PD genes. Number in parentheses indicates the number of expression features used in each case. **(B)** Correlation between screened Chronos score and predicted dependency score across 1121 cell lines. Each point represents a dependency score for one genetic dependency in one cell line. Pearson correlation coefficient was calculated; *P*-value computed from t-distribution. **(C)** Average performance (Pearson correlation coefficient between screened Chronos score and predicted dependency score) for 1298 DepOIs from Chiu et al. (*48*) using performance reported in Chiu et al. or using TrPLet (this study). **(D)** Shared PD genes (R≥0.2 between screened Chronos score and predicted dependency score during training) between Shi et al. (*47*) RNA only model (green), Shi et al. omics model (blue), and TrPLet (this study). **(E)** *t*-SNE projection based on RNA expression for tumors in TCGA and cell lines in the DepMap, restricted to common lineages. Tumors are colored based on annotated lineage and selected clusters are labeled. **(F)** Coefficient of determination (R^2^) between predicted dependency scores for tumors in TCGA and sample purity across 6,283 PD genes. **(G)** Predicted dependency scores for *CTNNB1* and *CDK4* across tumor types profiled in TCGA; tumors grouped by TCGA lineage and relative feature importance shown. Each point represents an individual feature (see **Methods**). Expression features associated with *CTNNB1* dependency included multiple β-catenin pathway members (e.g. *AXIN2*, *NKD1*, *ASCL2*, *BMP4*), while *CDK4* dependency included its interacting cyclin *CCND1* and paralog *CDK6*. **(H)** Relative feature importance for RNA predictors of *SOX10, FAM50A*, and *KRAS* dependency score. Each point represents an individual feature (see **Methods**). **(I)** Predicted dependency scores for *ERG* across TCGA prostate tumors (PRAD); tumors grouped by *ERG* fusion partner. *P*-values computed by Mann Whitney U-test; ****P<0.0001. **(J)** *Left*: Correlation between predicted *KRAS* dependency score and log_2_(fold-change) in cell viability upon single dose treatment (2.5 μM, 5 days) with the KRAS G12C inhibitor sotorasib in DepMap across non-G12C KRAS mutant cancer cell lines. *Right*: Correlation between experimentally-derived *KRAS* Chronos score (from CRISPR screens in DepMap) and log_2_(fold-change) in cell viability upon single dose treatment (2.5 μM, 5 days) with the KRAS G12C inhibitor sotorasib in DepMap across KRAS^G12C^ mutant cancer cell lines. Pearson correlation coefficient and *P*-value computed from t-distribution are annotated. **(K)** Correlation between predicted gene dependency score for PD genes and area under the curve (log_10_(concentration) vs. normalized cell viability) for selected compounds profiled in the DepMap. Pearson correlation coefficient and *P*-value computed from t-distribution are plotted for each combination. **(L)** Likelihood ratio test for skew-t vs. normal distribution for the distribution of each PD gene’s dependency score across TCGA (y-axis) vs. GTEx (x-axis).

**Fig. S2.**
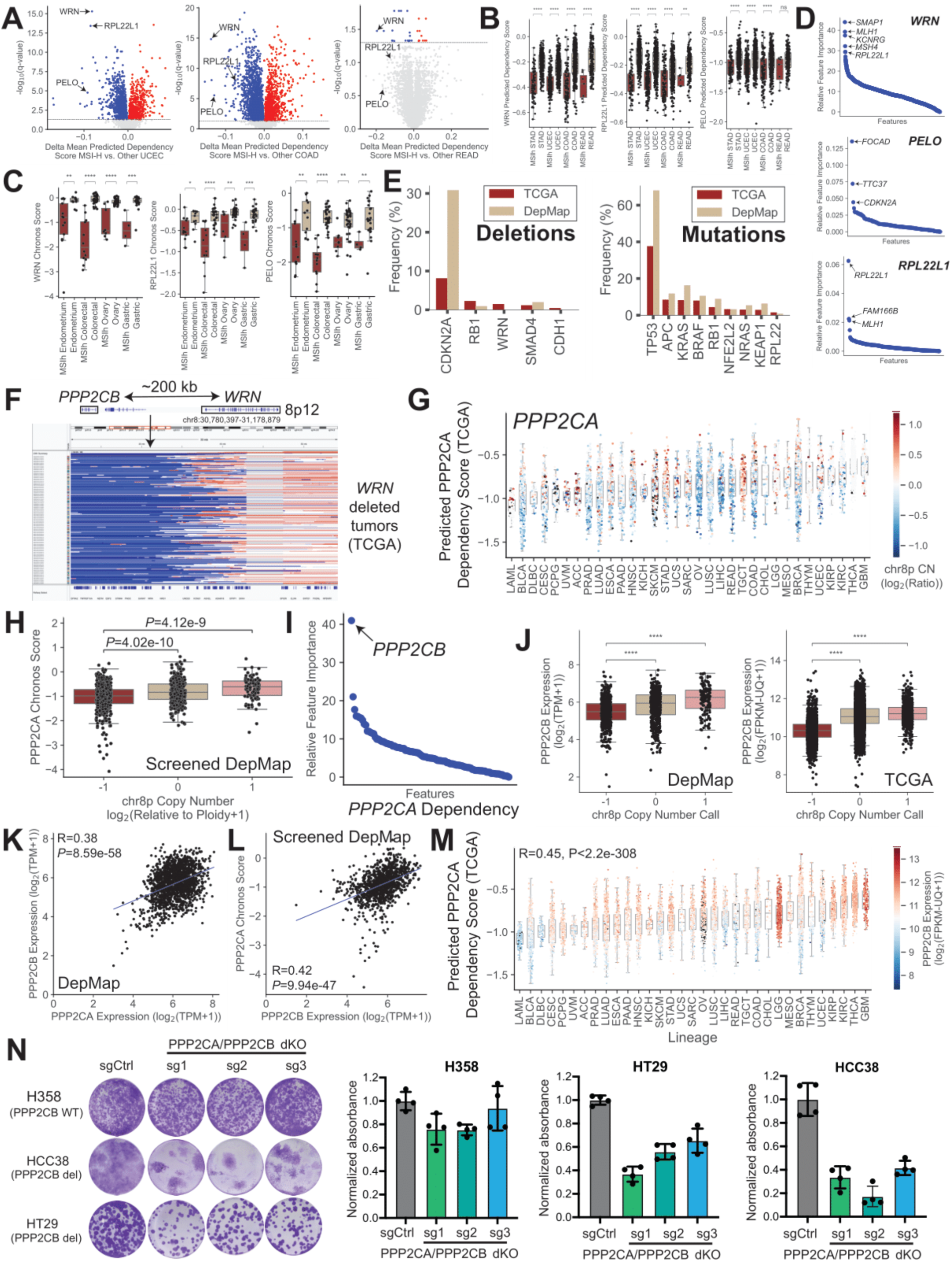
Virtual CRISPR screening uncovers synthetic lethal relationships from tumor transcriptomes. **(A)** Comparison of predicted dependencies in PD genes between MSI-H and MSS TCGA endometrial carcinomas (UCEC), colon adenocarcinomas (COAD), and rectal adenocarcinomas (READ). **(B)** Predicted dependency scores for *WRN*, *PELO*, and *RPL22L1* across TCGA stomach adenocarcinomas (STAD), endometrial carcinomas (UCEC), colon adenocarcinomas (COAD), and rectal adenocarcinomas (READ); tumors grouped by lineage and MSI status. *P*-values computed by Mann Whitney U-test; ****P<0.0001. **(C)** Chronos scores for *WRN*, *PELO*, and *RPL22L1* across screened gastric, endometrial, colorectal, and ovarian cancer cell lines in the DepMap; cell lines grouped by lineage and MSI status. *P*-values computed by Mann Whitney U-test; **P<0.01, ***P<0.001. **(D)** Relative feature importance for RNA predictors of *WRN, PELO*, and *RPL22L1* dependency score. Each point represents an individual feature (see **Methods**). **(E)** Comparison of alteration frequency between TCGA (red) and DepMap (brown) for deletions (left) and mutations (right) for genes shown in Fig. 2B**-C**. **(F)** IGV screenshot of all samples in TCGA with called WRN deletions (log_2_(copy-number/2) ≤ -1), demonstrating arm-level loss of chr8p in most cases. The zoomed in *WRN* locus is shown above the plot, with *PPP2CB* being located 200 kb from *WRN* and universally co-deleted. **(G)** Predicted dependency scores for *PPP2CA* across tumor types profiled in TCGA; tumors grouped by TCGA lineage. Samples are colored by chr8p copy number. **(H)** Boxplot of *PPP2CA* Chronos score across screened cell lines in the DepMap, grouped by chr8p copy number call (-1: deletion, 0: neutral, 1: gain). *P*-values computed by Mann-Whitney U test. **(I)** Ranked relative feature importance for RNA predictors of *PPP2CA* dependency score. Each point represents an individual feature (see **Methods**). **(J)** *PPP2CB* expression in DepMap cell lines (left) or TCGA tumors (right) grouped by chr8p copy number status (-1: deletion, 0: copy neutral, 1: gain) (*185*). *P*-values computed by Mann Whitney U-test; ****P<0.0001. **(K)** Correlation between expression of *PPP2CA* and *PPP2CB* in cell lines across the DepMap. **(L)** Correlation between expression of *PPP2CB* and *PPP2CA* Chronos score across screened cell lines in the DepMap. **(M)** Predicted dependency scores for *PPP2CA* across tumor types profiled in TCGA; tumors grouped by TCGA lineage. Samples are colored by *PPP2CB* expression. Pearson correlation coefficient was calculated; *P*-value computed from t-distribution. **(N)** Representative photograph (left) and quantification of colony formation assay (right) for *PPP2CB* WT (H358) and *PPP2CB* deleted (HCC38, HT29) cell lines after CRISPR/Cas9 double knockout of *PPP2CA/PPP2CB*. Shown as mean +/- s.d., n=4 biological replicates per condition.

**Fig. S3.**
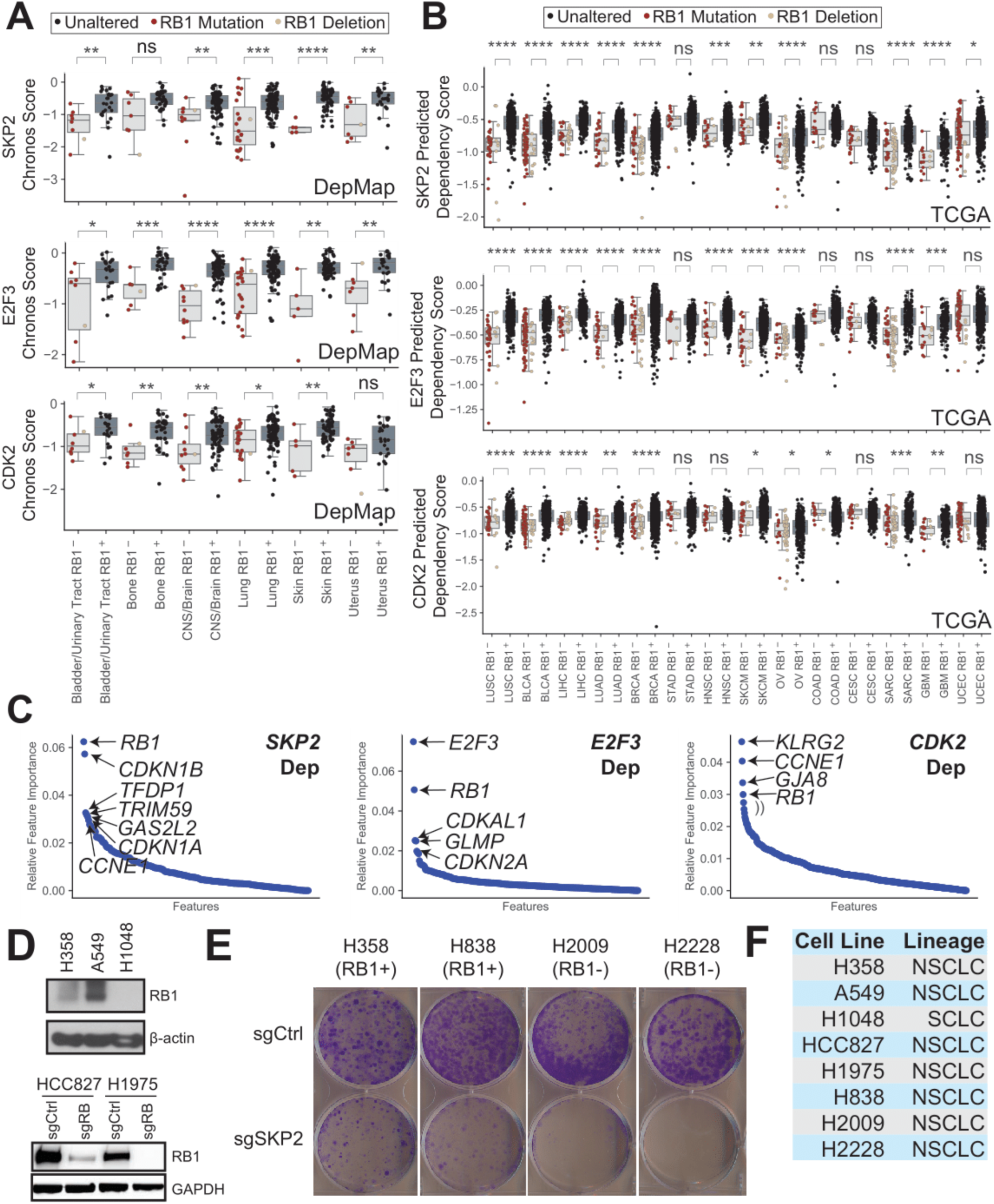
SKP2, E2F3, and CDK2 are synthetic lethal with Rb1 inactivation. **(A)** Chronos scores for *SKP2, E2F3,* and *CDK2* across screened DepMap cell lines in lineages with frequent *RB1* alteration; cell lines grouped by lineage and *RB1* loss-of-function status. Deletions and mutations are shown in the same plot. *P*-values computed by Mann Whitney U-test; n.s. not significant, *P<0.05, **P<0.01, ***P<0.001, ****P<0.0001. **(B)** Predicted dependency scores for *SKP2*, *E2F3*, and *CDK2* across TCGA in lineages with frequent *RB1* alteration; tumors grouped by lineage and *RB1* loss-of-function status. Deletions and mutations are shown in the same plot. *P*-values computed by Mann Whitney U-test; n.s. not significant, *P<0.05, **P<0.01, ***P<0.001, ****P<0.0001. **(C)** Relative feature importance for RNA predictors of *SKP2, E2F3*, and *CDK2* dependency score. Each point represents an individual feature (see **Methods**). **(D)** Western blot for RB1 and housekeeping gene (β-actin, GAPDH) in cell lines assayed in Fig. 2. **(E)** Representative colony formation assay for RB1+ (H358, H858) and RB1- (H2009, H2228) NSCLC cell lines after CRISPR/Cas9 knockout of *SKP2* (normalized to sgCtrl). Quantification of colony formation is shown in Fig. 2M. **(F)** DepMap cancer type (Oncotree Primary Disease) for cell lines assayed in Fig. 2.

**Fig. S4.**
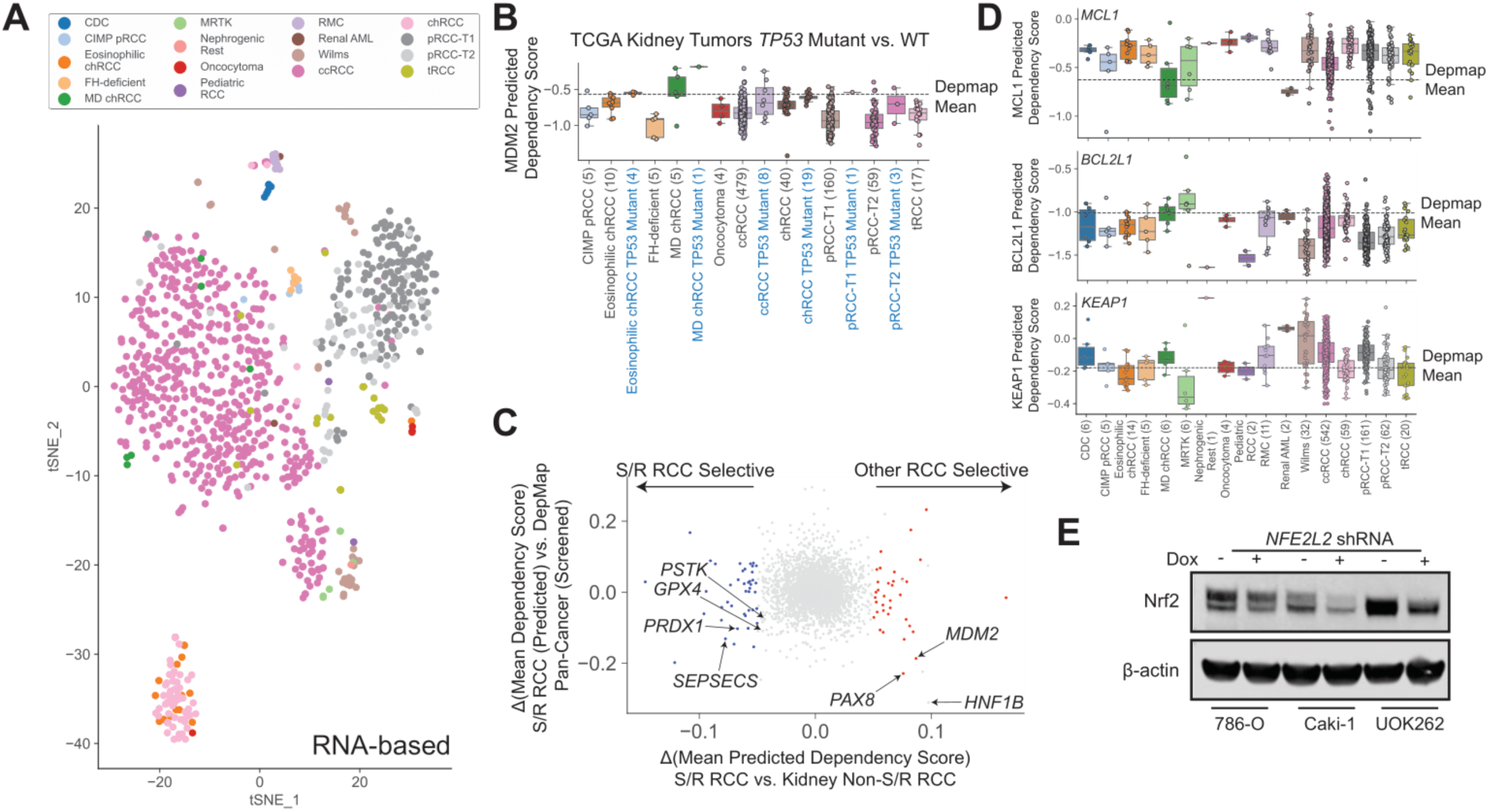
Virtual CRISPR screening predicts dependency landscape of rare kidney cancers. **(A)** *t*-SNE projection and hierarchical clustering based on RNA expression for kidney tumors with dependencies predicted by TrPLet in this manuscript (N=938), grouped by subtype as indicated in Fig. 3A. **(B)** Box plot of predicted *MDM2* dependency *score* across TCGA kidney cancers stratified by annotated *TP53* mutation status. DepMap mean Chronos score for *MDM2* is indicated as a dotted line. **(C)** Predicted dependency scores for S/R RCCs vs. non-S/R RCCs (ccRCC, pRCC-T1, pRCC-T2, chRCC) in TCGA. **(D)** Box plot of predicted *MCL1* (top), *BCL2L1* (middle), and *KEAP1* (bottom) dependency scores for individual kidney tumors (N=938), grouped by subtype indicated in Fig. 3A. Mean Chronos score in DepMap (experimentally derived) for these genes are indicated by dotted lines. **(E)** Validation of doxycycline-inducible knockdown of NRF2 expression by Western blotting.

**Fig. S5.**
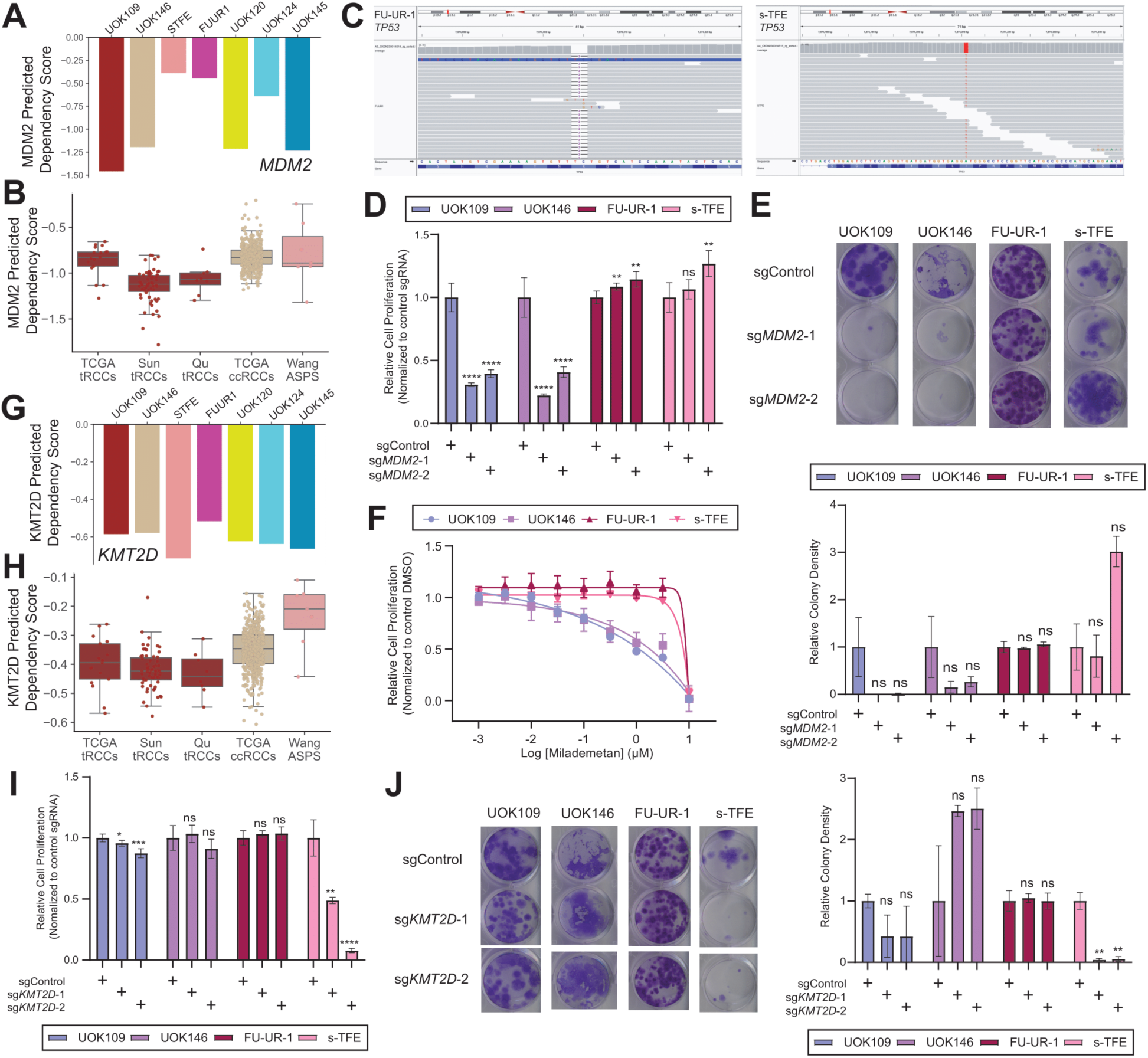
Validation of *MDM2* and *KMT2D* dependency in tRCC. **(A)** Predicted *MDM2* dependency score across 7 tRCC cell lines profiled by RNA-seq. UOK109, UOK146, UOK120, and UOK145 are predicted to be dependent on *MDM2*. **(B)** Predicted *MDM2* dependency score in either tRCC (TCGA, Sun et al., Qu et al.), ccRCC (TCGA), or ASPS (Wang et al.) RNA-seq tumor cohorts. Note, tRCC tumors have a higher frequency of predicted *MDM2* dependency than cell lines, likely due to acquisition of *TP53* mutations/expansion of *TP53* negative clones in cell culture. **(C)** IGV snapshots showing mutations at the TP53 locus in FU-UR-1 and s-TFE cells. **(D)** Relative viability of tRCC cells (UOK109, UOK146, FU-UR-1, s-TFE) after CRISPR/Cas9 knockout of *MDM2.* Viability is normalized to control sgRNA, shown as mean +/- s.d., n=6 biological replicates per condition. *P-*values were calculated by Welch’s (two-tailed unpaired) t-test as compared with sgControl samples. \**P* < 0.05, \*\**P* < 0.01, \*\*\**P* < 0.001, \*\*\*\**P* < 0.0001. UOK109 and UOK146 are sensitive; FU-UR-1 and S-TFE are resistant. **(E)** Representative photograph (left) and quantification of colony formation assay (right) for tRCC cells (UOK109, UOK146, FU-UR-1, s-TFE) after CRISPR/Cas9 knockout of *MDM2*. Shown as mean +/- s.d., n=3 biological replicates per condition. *P*-values were calculated by Welch’s (two-tailed unpaired) t-test as compared with sgControl samples. \**P* < 0.05, \*\**P* < 0.01, \*\*\**P* < 0.001, \*\*\*\**P* < 0.0001. **(F)** Viability of tRCC cell lines treated with indicated concentrations of milademetan (MDM2 inhibitor) and assayed for cell viability after 3 days with CellTiter-Glo. Viability at each concentration is relative to vehicle-treated cells, shown as mean +/- s.d., n=6 biological replicates. **(G)** Predicted *KMT2D* dependency score across 7 tRCC cell lines profiled by RNA-seq. S-TFE is predicted to be most dependent on *KMT2D*. **(H)** Predicted *KMT2D* dependency score in either tRCC (TCGA, Sun et al., Qu et al.), ccRCC (TCGA), or ASPS (Wang et al.) RNA-seq tumor cohorts. **(I)** Relative viability of tRCC cells (UOK109, UOK146, FU-UR-1, s-TFE) after CRISPR/Cas9 knockout of *KMT2D.* Viability is normalized to control sgRNA, shown as mean +/- s.d., n=6 biological replicates per condition. *P*-values were calculated by Welch’s (two-tailed unpaired) t-test as compared with sgControl samples. \**P* < 0.05, \*\**P* < 0.01, \*\*\**P* < 0.001, \*\*\*\**P* < 0.0001. **(J)** Representative photograph (left) and quantification of colony formation assay (right) for tRCC cells (UOK109, UOK146, FU-UR-1, s-TFE) after CRISPR/Cas9 knockout of *KMT2D*. Shown as mean +/- s.d., n=3 biological replicates per condition. *P*-values were calculated by Welch’s (two-tailed unpaired) t-test as compared with sgControl samples. \**P* < 0.05, \*\**P* < 0.01, \*\*\**P* < 0.001, \*\*\*\**P* < 0.0001.

**Fig. S6.**
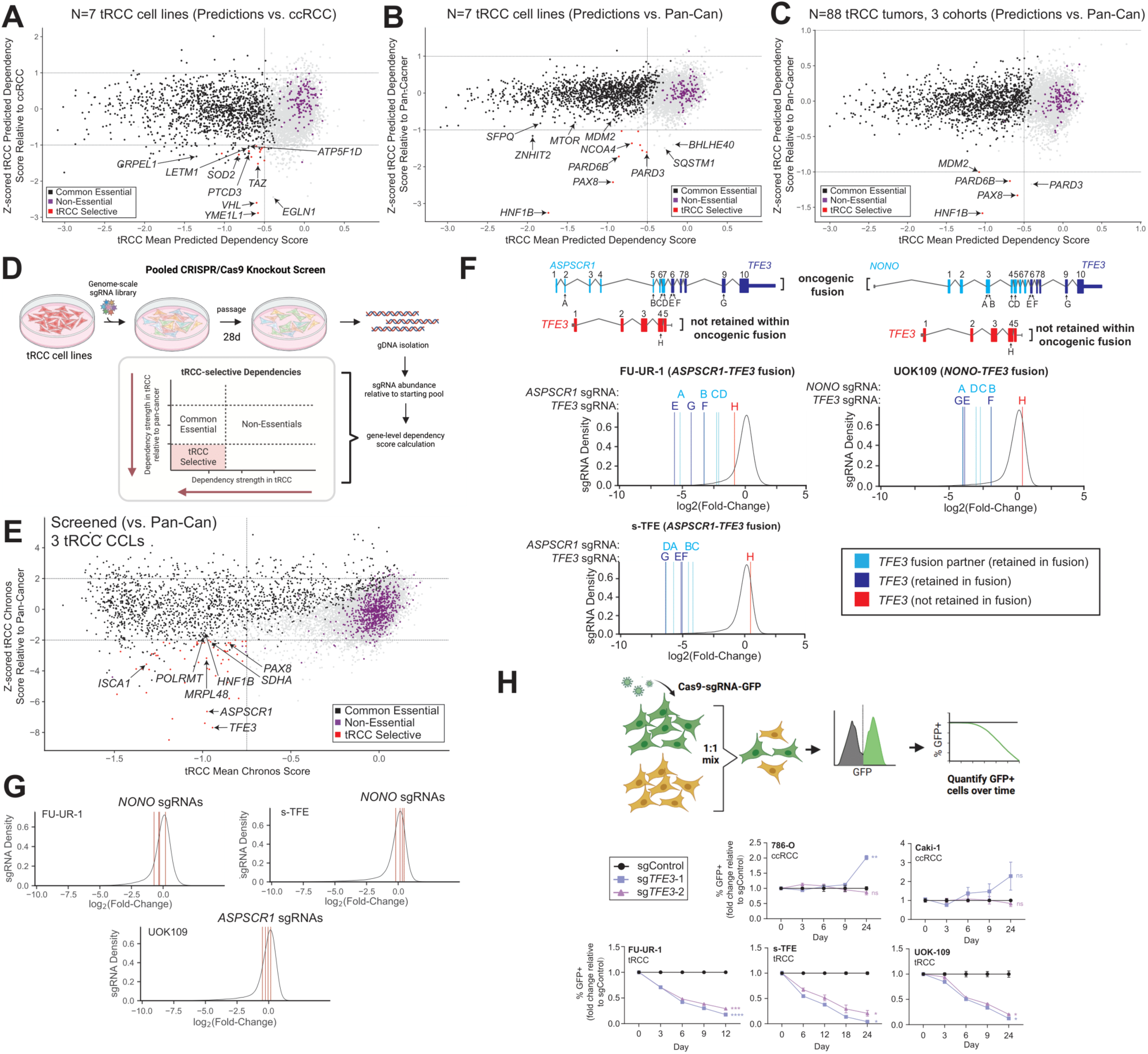
Genome-scale CRISPR screening reveals selective essentialities of tRCC cells. **(A-C)** Mean predicted dependency score for each PD gene across 7 tRCC cell lines **(A-B)** or 88 tRCC tumors from 3 cohorts **(C)** is plotted against the Z-scored predicted dependency score for that gene (relative to DepMap ccRCC cell lines **(A)** or DepMap pan-cancer **(B-C)**). Non-essential genes are colored purple while common essential genes are colored black (*184*). tRCC-selective dependencies are colored in red. **(D)** Workflow for CRISPR screens and analysis to identify tRCC-selective genetic dependencies. **(E)** Plot of tRCC-selective dependencies relative to all DepMap cell lines (pan-cancer). **(F)** Log-fold change for individual sgRNAs targeting either *TFE3* (E, F, G, H) or fusion partner (A, B, C, D) in tRCC CRISPR screens. The exons targeted by each sgRNA are indicated in the schematics. For each cell line, the top schematic represents the exons coding for the oncogenic fusion while the bottom schematic represents the N-terminal exons of *TFE3* not included in the oncogenic fusion. Density plot shows the distribution for LFC of all sgRNAs assessed in the CRISPR screen in each cell line while vertical lines represent log-fold change for individual sgRNAs. *Note*: Figure shows the *ASPSCR1-TFE3* fusion in s-TFE cells; the *ASPSCR1*-*TFE3* fusion in FU-UR-1 cells retains exon 5 of *TFE3* in the oncogenic fusion. **(G)** Log-fold change for individual sgRNAs targeting a fusion partner not involved in the translocation in each tRCC cell line (FU-UR-1 & S-TFE: *NONO*, UOK109: *ASPSCR1*), demonstrating non-essentiality. **(H)** Competitive growth assay to assess the effects of *TFE3* knockout in two ccRCC lines (786-O, Caki-1) and three tRCC lines (UOK109, FU-UR-1, s-TFE). Cells expressing Cas9/sgRNA and GFP were mixed in a 1:1 ratio with parental cells and proportion relative to sgControl cells was calculated at each time point via flow cytometry. Shown as mean +/- s.d., n=2 biological replicates per condition. *P*-values were calculated by Welch’s (two-tailed unpaired) t-test as compared with sgControl samples at the final time point. \**P* < 0.05, \*\**P* < 0.01, \*\*\**P* < 0.001, \*\*\*\**P* < 0.0001.

**Fig. S7.**
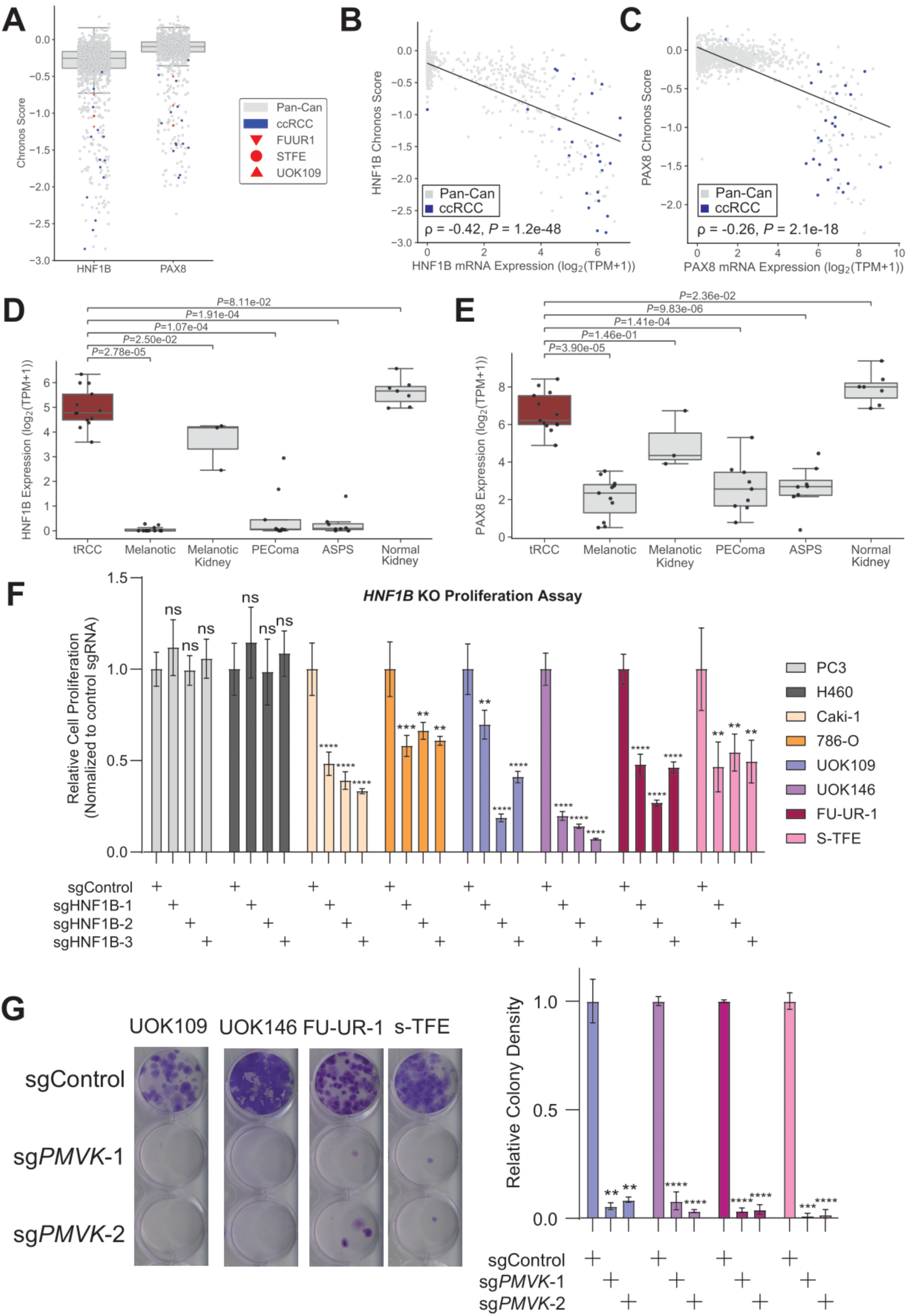
Validation of lineage dependencies *HNF1B* and *PMVK* in tRCC cells. **(A)** Distribution of Chronos scores for *HNF1B* and *PAX8* across DepMap cell lines and tRCC cell lines. ccRCC cell lines are shown in blue, tRCC cell lines are shown in red. **(B-C)** Correlation between *HNF1B* **(B)** or *PAX8* **(C)** Chronos score and gene expression (log_2_(TPM+1)) across DepMap cell lines. ccRCC cell lines are shown in blue. *P*-values computed by Spearman’s rho test. **(D-E)** In a published RNA-seq dataset (*126*), the expression of *HNF1B* **(D)** and *PAX8* **(E)** are shown (log_2_(TPM+1)) across kidney-derived *TFE3* fusion tumors (tRCC, melanotic kidney), normal kidney tissue, and other TFE3-driven tumors (PEComa, ASPS, Melanotic). **(F)** Viability of nonRCC cells (PC3, H460), ccRCC cells (Caki-1 and 786-O) or tRCC cells (UOK109, UOK146, FU-UR-1, s-TFE) after *HNF1B* knockout. Viability was measured by CellTiter-Glo; n=6 replicates per condition. *P*-values were calculated by Welch’s (two-tailed unpaired) t-test as compared with sgControl samples. \**P* < 0.05, \*\**P* < 0.01, \*\*\**P* < 0.001, \*\*\*\**P* < 0.0001. **(G)** Representative photograph (left) and quantification of colony formation assay (right) for tRCC cells (UOK109, UOK146, FU-UR-1, s-TFE) after CRISPR/Cas9 knockout of *PMVK*. Shown as mean +/- s.d., n=3 biological replicates per condition. *P*-values were calculated by Welch’s (two-tailed unpaired) t-test as compared with sgControl samples. \**P* < 0.05, \*\**P* < 0.01, \*\*\**P* < 0.001, \*\*\*\**P* < 0.0001.

**Fig. S8.**
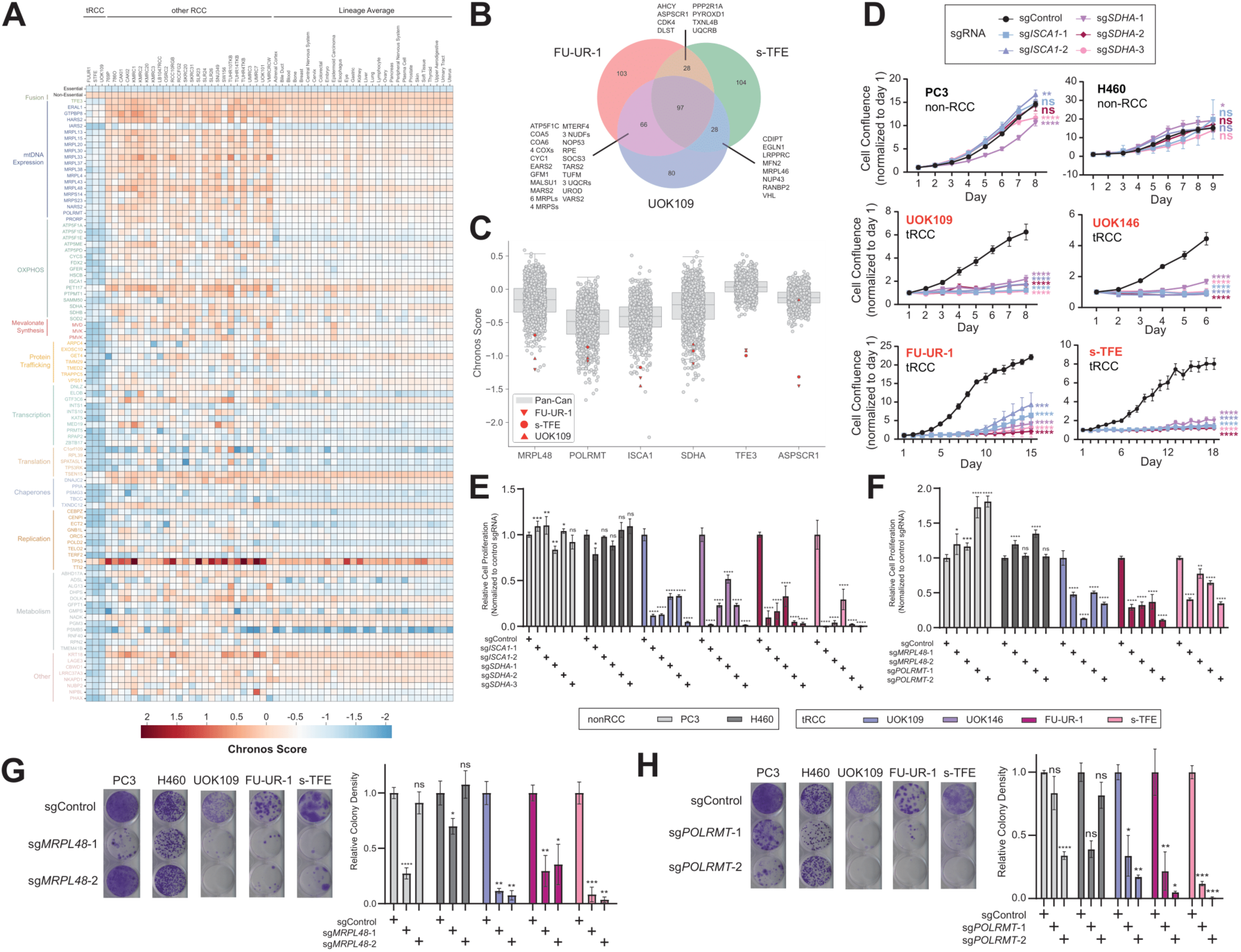
Predicting and validating mitochondrial and oxidative phosphorylation dependencies in tRCC. **(A)** Heat map of tRCC-selective dependencies (defined as genes with ΔChronos ≤ -0.5 between every tRCC cell line and DepMap ccRCC mean) grouped by pathway. Chronos scores for individual tRCC cell lines, DepMap RCC cell lines, and average Chronos score for each of 28 lineages screened in DepMap are shown. The top two rows indicate mean Chronos scores for essential and non-essential genes in each column, shown for reference. **(B)** Overlapping dependencies between tRCC cell lines screened in this study (gene included if ΔChronos ≤ -0.5 between an individual cell line and DepMap ccRCC mean). **(C)** Distribution of Chronos scores for indicated genes (*POLRMT*, *MRPL48*, *ISCA1, SDHA, TFE3, ASPSCR1*) across all DepMap cell lines (gray) and tRCC cell lines screened in this study (red). **(D)** Relative confluence of non-RCC cells (PC3, H460), and tRCC cells (UOK109, UOK146, FU-UR-1, s-TFE) after infection with lentivirus expressing Cas9 and either non-targeting control sgRNA, *ISCA1* sgRNAs, or *SDHA* sgRNAs. Confluence was normalized to day 1, shown as mean +/- s.d., n=6 biological replicates per condition. *P*-values were calculated by Welch’s (two-tailed unpaired) t test as compared with sgControl samples for the last assay day. \**P* < 0.05, \*\**P* < 0.01, \*\*\**P* < 0.001, \*\*\*\**P* < 0.0001. **(E-F)** Relative viability of non-RCC cells (PC3, H460) or tRCC cells (UOK109, UOK146, FU-UR-1, s-TFE) after CRISPR/Cas9 knockout of *ISCA1* **(E)**, *SDHA* **(E)**, *MRPL48* **(F)** or *POLRMT* **(F).** Viability is normalized to control sgRNA, shown as mean +/- s.d., n=6 biological replicates per condition. *P*-values were calculated by Welch’s (two-tailed unpaired) t-test as compared with sgControl samples. \**P* < 0.05, \*\**P* < 0.01, \*\*\**P* < 0.001, \*\*\*\**P* < 0.0001. **(G-H)** Representative photograph (left) and quantification of colony formation assay (right) for of non-RCC cells (PC3, H460) or tRCC cells (UOK109, FU-UR-1, s-TFE) after CRISPR/Cas9 knockout of *MRPL48* **(G)** or *POLRMT* **(H)**. Shown as mean +/- s.d., n=3 biological replicates per condition. *P*-values were calculated by Welch’s (two-tailed unpaired) t-test as compared with sgControl samples. \**P* < 0.05, \*\**P* < 0.01, \*\*\**P* < 0.001, \*\*\*\**P* < 0.0001.

**Fig. S9.**
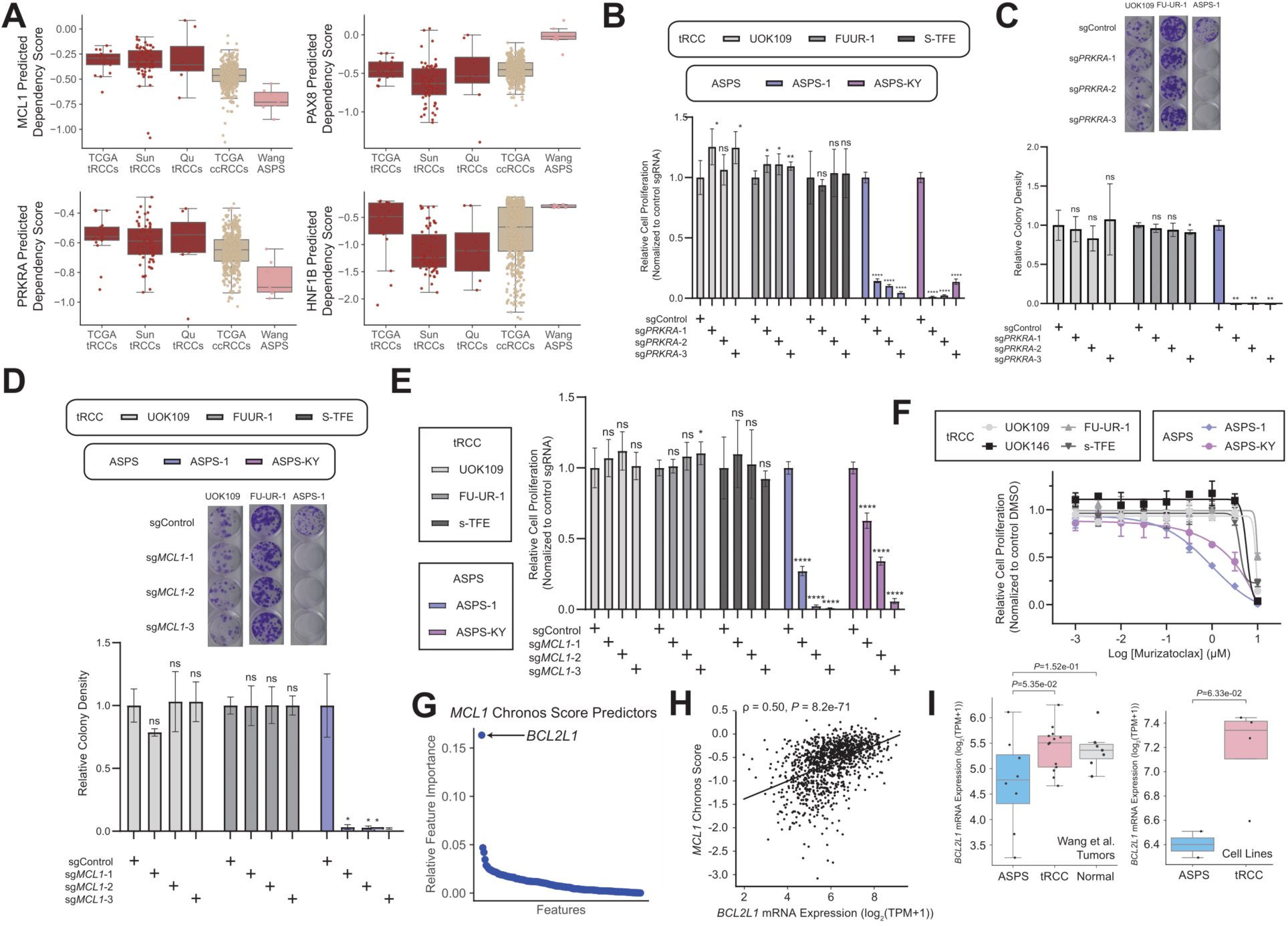
Predicting and validating *PRKRA* and *MCL1* dependency in ASPS. **(A)** Predicted dependency scores from tumor samples in either tRCC (TCGA, Sun et al., Qu et al.), ccRCC (TCGA), or ASPS (Wang et al.) RNA-Seq cohorts for the following genes: *MCL1*, *PRKRA*, *PAX8*, *HNF1B*. **(B)** Proliferation of tRCC cells (UOK109, FU-UR-1, s-TFE) and ASPS cell lines (ASPS-1 and ASPS-KY) transduced with one of 3 distinct sgRNAs targeting *PRKRA* or a non-targeting sgRNA control. n = 6 biological replicates per condition. *P*-values were calculated by Welch’s (two-tailed unpaired) t-test as compared with sgControl samples. \**P* < 0.05, \*\**P* < 0.01, \*\*\**P* < 0.001, \*\*\*\**P* < 0.0001. **(C-D)** Representative photograph (top) and quantification of colony formation assay (bottom) for of tRCC cells (UOK109, FU-UR-1) or ASPS cells (ASPS-1) after CRISPR/Cas9 knockout of *PRKRA* **(C)** and *MCL1* **(D)**. Shown as mean +/- s.d., n=3 biological replicates per condition. *P*-value were calculated by Welch’s (two-tailed unpaired) t-test as compared with sgControl samples. \**P* < 0.05, \*\**P* < 0.01, \*\*\**P* < 0.001, \*\*\*\**P* < 0.0001. **(E)** Proliferation of ASPS cell lines transduced with one of 3 distinct sgRNAs targeting *MCL1* or a non-targeting sgRNA control. Shown as mean +/- s.d., n = 6 biological replicates per condition. *P*-values were calculated by Welch’s t-test (two-tailed unpaired) as compared with sgControl samples. \**P* < 0.05, \*\**P* < 0.01, \*\*\**P* < 0.001, \*\*\*\**P* < 0.0001. **(F)** Viability of ASPS-1 and ASPS-KY and non-ASPS (versus tRCC) cell lines treated with indicated concentrations of murizatoclax (MCL1 inhibitor) and assayed for cell viability after 3 days with CellTiter-Glo. Viability at each concentration is relative to vehicle-treated cells, shown as mean +/- s.d., n=3 replicates. **(G)** Relative feature importance (ranked across top 5000 features) for RNA predictors of *MCL1* dependency score. Each point represents an individual feature (see **Methods**). **(H)** *MCL1* Chronos score plotted against *BCL2L1* mRNA expression (log_2_(TPM+1)) across all DepMap cell lines. **(I)** *BCL2L1* mRNA expression (log_2_(TPM+1)) in ASPS tumors, tRCC tumors, and kidney-adjacent normal tissue from Wang et al. (*126*) (left) as well as ASPS cell lines (ASPS-1, ASPS-KY) and tRCC cell lines (FU-UR-1, s-TFE, UOK109, UOK146) (right) profiled by RNA-seq are shown. *P*-values computed by Welch’s (two-tailed unpaired) t-test.

